# microRNA-1 Regulates Metabolic Flexibility in Skeletal Muscle via Pyruvate Metabolism

**DOI:** 10.1101/2024.08.09.607377

**Authors:** Ahmed Ismaeel, Bailey D. Peck, McLane M. Montgomery, Benjamin I. Burke, Jensen Goh, Gyumin Kang, Abigail B. Franco, Qin Xia, Katarzyna Goljanek-Whysall, Brian McDonagh, Jared M. McLendon, Pieter J. Koopmans, Daniel Jacko, Kirill Schaaf, Wilhelm Bloch, Sebastian Gehlert, Yuan Wen, Kevin A. Murach, Charlotte A. Peterson, Ryan L. Boudreau, Kelsey H. Fisher-Wellman, John J. McCarthy

## Abstract

MicroRNA-1 (miR-1) is the most abundant miRNA in adult skeletal muscle. To determine the function of miR-1 in adult skeletal muscle, we generated an inducible, skeletal muscle-specific miR-1 knockout (KO) mouse. Integration of RNA-sequencing (RNA-seq) data from miR-1 KO muscle with Argonaute 2 enhanced crosslinking and immunoprecipitation sequencing (AGO2 eCLIP-seq) from human skeletal muscle identified miR-1 target genes involved with glycolysis and pyruvate metabolism. The loss of miR-1 in skeletal muscle induced cancer-like metabolic reprogramming, as shown by higher pyruvate kinase muscle isozyme M2 (PKM2) protein levels, which promoted glycolysis. Comprehensive bioenergetic and metabolic phenotyping combined with skeletal muscle proteomics and metabolomics further demonstrated that miR-1 KO induced metabolic inflexibility as a result of pyruvate oxidation resistance. While the genetic loss of miR-1 reduced endurance exercise performance in mice and in *C. elegans,* the physiological down-regulation of miR-1 expression in response to a hypertrophic stimulus in both humans and mice causes a similar metabolic reprogramming that supports muscle cell growth. Taken together, these data identify a novel post-translational mechanism of adult skeletal muscle metabolism regulation mediated by miR-1.

## Introduction

Skeletal muscle is the largest organ in the body, comprising about 35% of the human body by mass [1]. Skeletal muscle mechanical activity is required for posture, movement, and breathing [2]. The loss of muscle mass associated with aging and wasting diseases such as cancer cachexia results in increased dependence, frailty, and mortality, and reduced quality of life [3]. Skeletal muscle is also responsible for 70-80% of postprandial or insulin-stimulated glucose uptake from circulation. This tissue is therefore recognized as having a central role in whole-body metabolism, influencing metabolic diseases such as diabetes and obesity [4]. A better understanding of the factors regulating skeletal muscle mass and metabolism is critical for the development of effective therapies to maintain and/or restore muscle function with age and disease.

Almost a decade after the first description of microRNAs (miRNAs) as key mediators of post-transcriptional gene regulation [5, 6], research began to reveal cell- and tissue-specific enrichment of different sets of miRNAs [7, 8]. microRNA-1 (miR-1) was one of the first identified miRNAs [9], and among the first miRNAs found to be enriched in particular organs (heart and skeletal muscle for miR-1) [10]. Of the striated muscle-enriched miRNAs (myomiRs), miR-1 is by far the most abundant miRNA in adult human, rodent, equine, ovine, caprine, bovine, and porcine skeletal muscle, accounting in some studies for up to 80-90% of all miRNA reads [11–17].

A unique aspect of miR-1 is its degree of conservation as one of only 32 miRs conserved among all Bilaterian animals [18]. The conservation of miR-1 encompasses both its sequence as well as its muscle-specific enrichment, suggesting an integral role in regulating muscle function [19]. For example, miR-1 of *Drosophila* differs by only a single nucleotide from *C. elegans* and human miR-1 [20]. Although miR-1 has been shown to be critical for proper muscle development in both invertebrates and vertebrates using *in vitro* techniques [21–34], technical barriers have prevented further interrogation of how miR-1 controls myogenesis *in vivo* [35].

Furthermore, the specific function of miR-1 in adult skeletal muscle remains to be rigorously investigated. Instead, previous studies using knockout (KO) approaches to study miR-1 have all used germline models or young, developing mice [36–38]. These *in vivo* studies also involved deletion of the other myomiRs along with miR-1, including miR-133 and miR-206, preventing a clear delineation of the specific function of miR-1 [36–38].

A major challenge in determining the function of miR-1 in adult skeletal muscle is the rigorous identification of bona fide target genes. The typical *in silico* approaches overpredict miRNA binding sites, inaccurately model target site accessibility, and miss biologically significant miRNA binding [39–41]. Furthermore, popular *in vitro* methods such as luciferase reporter assays do not account for *in vivo* stochiometric relationships between miRNAs and accessible mRNA targets in the cell type of interest, and therefore do not accurately reflect true biological interactions *in vivo* [39–41].

To address the aforementioned challenges, we used Argonaute 2 enhanced crosslinking and immunoprecipitation sequencing (AGO2 eCLIP-seq) data generated from the first transcriptome-wide map of bona fide miRNA target genes in human skeletal muscle tissue. We also generated an inducible mouse model to specifically inactivate only miR-1 in adult skeletal muscle without affecting other myomiRs. RNA-sequencing (RNA-seq), comprehensive mitochondrial and metabolic phenotyping, and metabolomics analyses revealed altered pyruvate metabolism in miR-1 KO mice. Integration of RNA-seq and eCLIP-seq datasets identified several biologically relevant miR-1 target genes that modulate pyruvate metabolism, including pyruvate kinase muscle isozyme (*Pkm*) and polypyrimidine tract binding protein (*Ptbp1*). PTBP1 mediates the alternative splicing of the *Pkm* gene in favor of the *Pkm2* isoform which promotes aerobic glycolysis by shuttling pyruvate away from the TCA cycle, thereby causing metabolic inflexibility. While the genetic loss of miR-1 causes a state of metabolic inflexibility that significantly reduces endurance exercise performance, we provide evidence that the physiological down-regulation of miR-1 in response to a hypertrophic stimulus in both humans and mice causes a similar metabolic reprogramming that may support muscle cell growth. In addition to elucidating a novel function for miR-1 in adult skeletal muscle, our integrated analysis of eCLIP-seq data along with a miRNA KO mouse model provides a highly efficacious framework for future *in vivo* miRNA interrogations in skeletal muscle homeostasis.

## Results

### miR-1 exhibits widespread binding in human skeletal muscle

The gold standard for identifying miRNA target genes is AGO2 eCLIP-seq because it allows for the direct biochemical determination of AGO2:mRNA binding. As part of a parallel initiative to generate an atlas of empirically defined miRNA target binding sites across human tissues, we performed AGO2 eCLIP on six human skeletal muscle tissue samples to generate a high-quality map of miRNA binding sites across the muscle transcriptome. We then leveraged empirical binding events (defined as significantly enriched peaks known as clusters, FDR < 0.05) with target site prediction simulation tools (TargetScan8.0 and miRcode) to identify 1286 miR-1 target sites across 1234 clusters and 1136 genes (Figure 1A). As miR-1 and miR-206 share identical seed sequences [42], we could not distinguish miR-1 sites from miR-206 sites.

**Figure 1.**
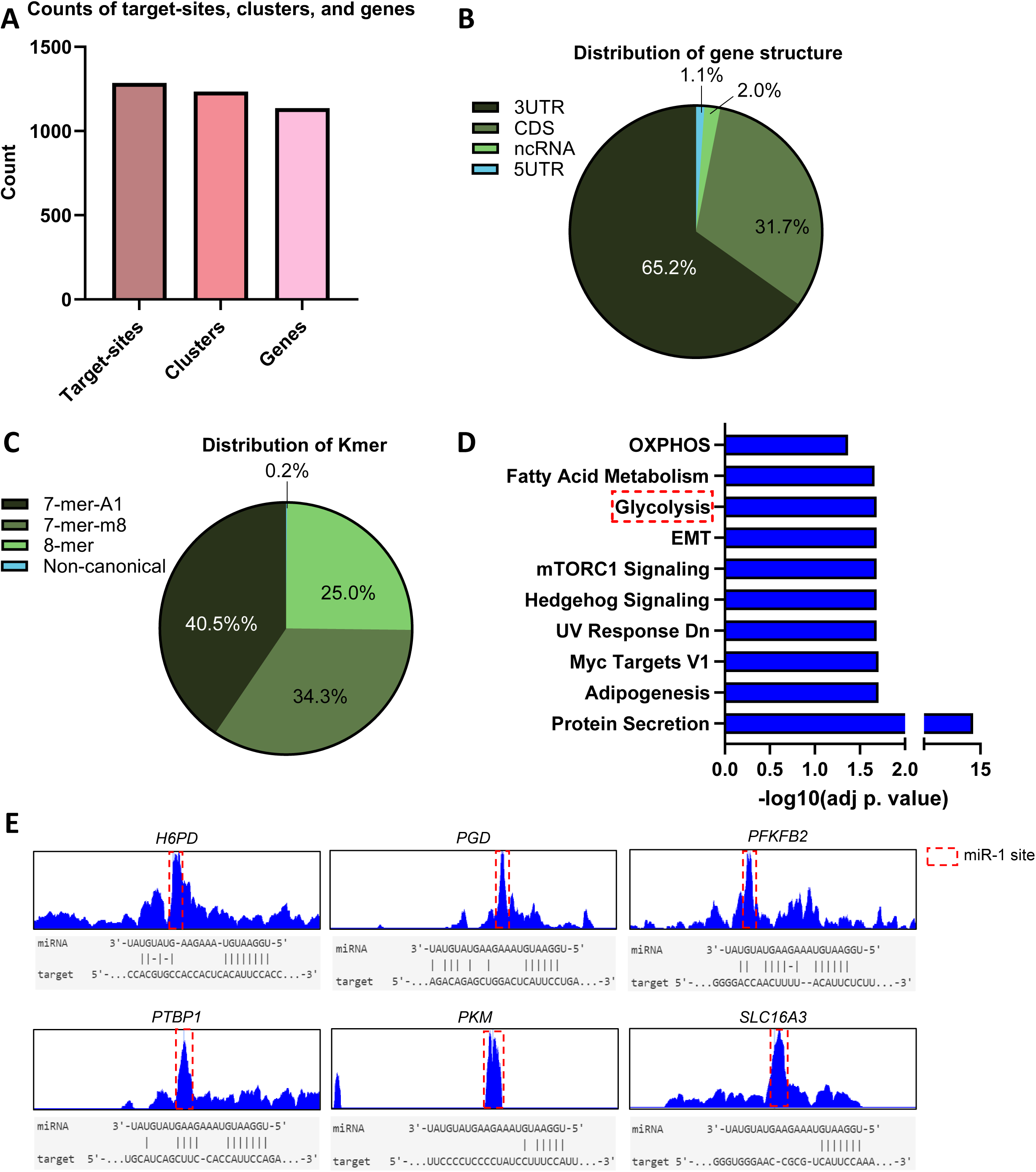
Widespread binding of miR-1 targets in human skeletal muscle. **(A)** Number of miR-1 target sites across clusters and genes (Count). **(B)** Pie chart of miR-1 binding events across gene structure, proportion (%) indicated. 3UTR: 3’ untranslated region, CDS: coding sequence, ncRNA: noncoding RNA, 5UTR: 5’ untranslated region. **(C)** Pie chart of Kmer distributions of miR-1 predicted binding events, proportion (%) indicated. **(D)** Pathway enrichment analysis (MSigDB Hallmark 2020) of AGO2 eCLIP-defined miR-1 target genes. Top 10 pathways by adj. *p*-value shown. **(E)** Example miR-1 binding peaks in mRNAs involved in glycolysis and parallel metabolic pathways outlined with red dotted line. miR-1 alignment on mouse target sequence shown below.

However, miR-1 is significantly (∼250-fold) more abundant in adult skeletal muscle than miR-206 [12]. In addition, miR-206 is more enriched in muscle stem cells (MuSCs) [43–45], so for clarity, we hereafter refer to the miR-1/206 seed sequence as just miR-1. Binding-events across gene structure demonstrate a preference 3UTR>CDS>ncRNA>5UTR (Figure1B). Kmer distributions of miR-1 predicted binding events show a broad distribution among canonical binding events 7merA1 (40.5%), 7mer8 (34.3%) 8mer (25%), non-canonical (0.2%) (Figure 1C).

Pathway analysis of eCLIP-defined miR-1 targets identified enrichment of protein secretory pathways, signaling pathways (Hedgehog, mammalian target of rapamycin complex 1, mTORC1), and metabolic pathways (glycolysis, fatty acid metabolism, and oxidative phosphorylation, Figure 1D). Of note, several miR-1 target genes were related to glycolysis, including hexose 6-phosphate dehydrogenase (*H6PD*), 6-phosphogluconate dehydrogenase (*PGD*), 6-phosphofructo-2-kinase (*PFKFB2*), polypyrimidine tract-binding protein 1 (*PTBP1*), pyruvate kinase muscle (*PKM*), and monocarboxylic acid transporter 4 (*SLC16A3*), with ScanMiR [46] confirming miR-1 binding sites in the orthologous mouse genes (Figure 1E).

### Generation of inducible skeletal muscle-specific miR-1 KO mouse

A major challenge for determining the specific function of miR-1 in adult skeletal muscle is the fact that the mature miR-1 is derived from two separate genes that encode two distinct bicistronic primary miRNA transcripts that contain miR-1-1 and miR-1-2, and miR-133a2 and miR-133a1, respectively. The miRNA-1-2 cluster resides within the intron of the Mind Bomb 1 (*Mib1*) gene [47]. Thus, while previous studies have used germline miR-1/133 knockouts [36], we sought to generate a mouse model for specific, inducible deletion of miR-1 in adult skeletal muscle, without affecting nearby miR-133 or *Mib1* expression, to determine the specific function of miR-1 *in vivo*. To this end, we crossed the skeletal muscle-specific inducible *Cre* mouse (HSA-MCM) [48] with a floxed *miR-1-1* and floxed *miR-1-2* mouse (*miR-1-1^f/f^; miR-1-2^f/f^*) [47] to generate the HSA-MCM; *miR-1-1^f/f^; miR-1-2^f/f^* mouse, designated HSA-miR-1 (Supplemental Figure S1A). Tamoxifen-treated littermate *miR-1-1^f/f^; miR-1-2^f/f^* mice (negative for HSA-MCM) or vehicle-treated HSA-miR-1 mice served as controls (wild-type, WT).

At 4 months of age, HSA-miR-1 mice were treated with tamoxifen to induce inactivation of miR-1-1 and miR-1-2 (miR-1 KO). After an 8-week washout period, whole-muscle miR-1 expression was ∼60 and ∼70% lower in soleus and plantaris muscles of miR-1 KO mice, respectively, compared to WT mice (Supplemental Figures S1B-C). However, in isolated single myofibers, the expression of miR-1 was reduced by >90% in miR-1 KO fibers (Supplemental Figure S1D). Importantly, skeletal muscle expression of miR-133a, *Mib1*, and miR-206 were not affected in miR-1 KO mice, nor was cardiac miR-1 expression, as assessed by quantitative PCR (qPCR) (Supplemental Figures S1E-H). Thus, phenotypes of miR-1 KO mice are attributed to the specific loss of skeletal muscle miR-1.

### Loss of miR-1 in skeletal muscle results in a cancer-like metabolic reprogramming gene expression signature

We next sought to assess the effect of miR-1 deletion on the transcriptome in adult skeletal muscle. To this end, we conducted RNA-seq of total RNA from gastrocnemius muscles of WT (n=4) and miR-1 KO (n=4) mice. Analysis of differentially up-regulated genes in miR-1 KO demonstrated enrichment of glycolytic pathways relative to WT (i.e., glycolysis, metabolic reprogramming in cancer, Warburg effect) as well as growth-related pathways (i.e., mTOR signaling, myostatin-IGF1 crosstalk) (Figure 2A). On the other hand, down-regulated genes in miR-1 KO included fatty acid oxidation pathways (i.e., mitochondrial beta oxidation, fatty acid beta oxidation, PPAR signaling pathway) (Figures 2B-C).

**Figure 2.**
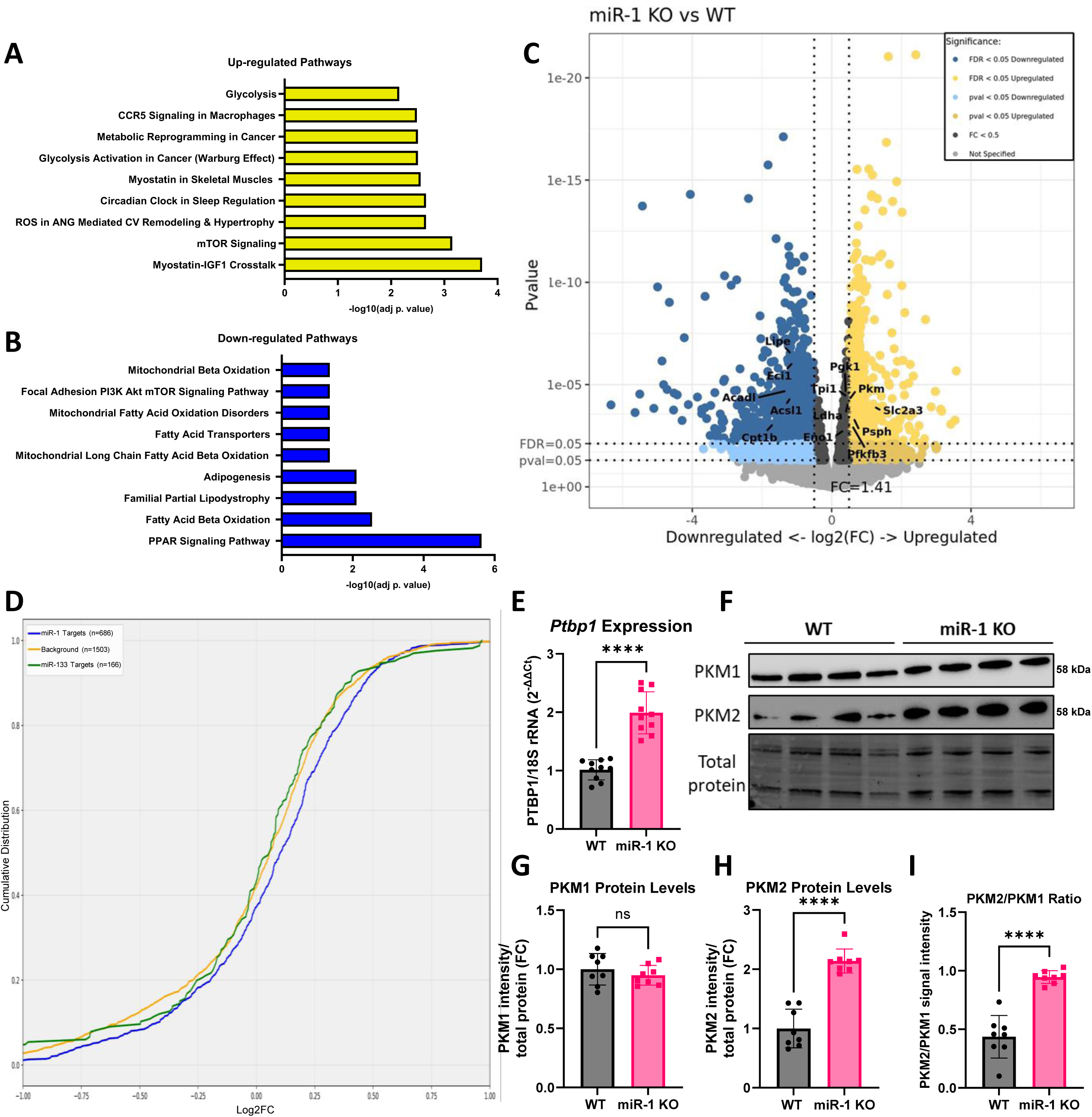
Transcriptomic profiling of miR-1 KO skeletal muscle. **(A-B)** Pathway enrichment analysis (Elsevier Pathway Collection) of significantly up-regulated genes **(A)** (FDR < 0.05, Log2FC > 0) and significantly down-regulated genes **(B)** (FDR < 0.05, Log2FC < 0) in miR-1 KO/WT from gastrocnemius bulk RNA-seq (n=4 female WT, n=4 female miR-1 KO, post-vehicle or tamoxifen treatment and an 8-week washout). Top 9 pathways (by adj. *p*-value) shown. **(C)** Volcano plot of differentially expressed genes, dashed lines indicate *p*-value, FDR, and FC cutoffs. Yellow represents significantly up-regulated genes, and blue represents significantly down-regulated genes. *P*-value cutoff set at 1e-25 for visibility. Genes in pathways from **(A-B)** labeled on the volcano plot. **(D)** Cumulative density function (CDF) of eCLIP-seq-defined miR-1 targets (blue) and miR-133 (green) Log2FC from RNA-seq compared to non-miR-1 targets (yellow). **(E)** *Ptbp1* mRNA expression in WT and miR-1 KO gastrocnemius. **(F)** Representative Western blot of PKM1 and PKM2 and corresponding total protein levels in WT and miR-1 KO gastrocnemius muscle lysates. **(G-I)** Quantification of **(G)** PKM1 protein levels, **(H)** PKM2 protein levels, and **(I)** the ratio of PKM2 to PKM1. For Western blotting and qPCR experiments, n=8-10 WT and n=8-10 miR-1 KO females (post-tamoxifen treatment and an 8-week washout) were used, and differences between WT and miR-1 KO were tested using an independent *t*-test. \*\*\*\**p* < 0.0001, ns: not significant.

Despite the up-regulation of growth-related pathways in miR-1 KO, loss of miR-1 for 8 weeks did not affect the muscle fiber size of fast-twitch, glycolytic plantaris or slow-twitch, oxidative soleus muscles (Supplemental Figures S2A-B). There was also no difference in fiber-type-specific cross-sectional area (CSA) in plantaris or soleus fibers (Supplemental Figures S2C-J). Moreover, there were only nominal differences in fiber-type composition between miR-1 KO and WT plantaris and soleus muscles which pointed towards a minor transition towards more fast-twitch, glycolytic fibers in the KO mice (Supplemental Figures S2K-L).

### Identification of functional miR-1 target-site response elements in adult mouse skeletal muscle

Next, we used a curated list of high confidence miR-1 target-genes from the Human AGO2 eCLIP to stratify expression perturbations following miR-1 KO. Cumulative distribution function (CDF) plots demonstrate that 3’UTR eCLIP-supported miR-1 targets were globally up-regulated in miR-1 KO muscles (as indicated by a rightward shift, *p* = 7.71e-4, KS = 0.0909 relative to non-miR-1 targets, Figure 2D). Parallel analysis of target genes of miR-133abc (the second most abundant miRNA in skeletal muscle) did not show any enhanced repression in the absence of miR-1 (Figure 2D, *p* = 0.749, KS = 0.0541).

To determine which miR-1 target genes have been previously shown to affect metabolic processes, the list of eCLIP-seq-defined miR-1 target genes was analyzed using the Rummagene tool, which finds matching gene sets in the literature [49]. Gene sets identified by Rummagene revealed a relationship between these miR-1 targets and previous studies reporting higher expression of the splicing factor PTBP1 [50–56]. Collectively, these studies found that the elevated expression of PTBP1 promotes the replacement of exon 9 for exon 10 in the pyruvate kinase muscle (*PKM*) mRNA to generate the PKM2 isoform [57–62] and that dominant expression of PKM2 over PKM1 maintains the Warburg effect in cells [57–62]. Up-regulated *Ptbp1* expression in miR-1 KO gastrocnemius muscle was confirmed by qPCR analysis (Figure 2E). PKM2 protein levels were also significantly higher in miR-1 KO gastrocnemius muscle, resulting in a higher ratio of PKM2 to PKM1 (Figures 2F-I). Thus, miR-1 directly regulates the alternative splicing and expression of PKM by targeting both *Ptbp1* and *Pkm* genes. Notably, RNA-seq analyses identified up-regulation of several other glycolytic and pentose phosphate pathway genes in miR-1 KO muscle as well (Supplemental Figures 3A-K), including genes not directly targeted by miR-1. Taken together, these data suggest that miR-1 directly and indirectly regulates skeletal muscle glycolytic gene expression.

### Bioenergetic phenotyping of miR-1 KO mitochondria identifies metabolic inflexibility and altered pyruvate metabolism

To investigate whether the changes in the mRNA/protein levels of PTBP1 and PKM1/2 associate with skeletal muscle bioenergetics, we performed high-resolution respirometry (HRR) to assess mitochondrial respiration of permeabilized soleus fibers. Pyruvate/malate (Pyr/Mal)-supported LEAK and ADP-stimulated respiration were significantly lower in the miR-1 KO compared to WT (Figure 3A). Furthermore, maximum respiration (FCCP-stimulated) was significantly lower in the miR-1 KO; however, there was no difference in complex II (succinate-supported) respiration between miR-1 KO and WT soleus muscles.

**Figure 3.**
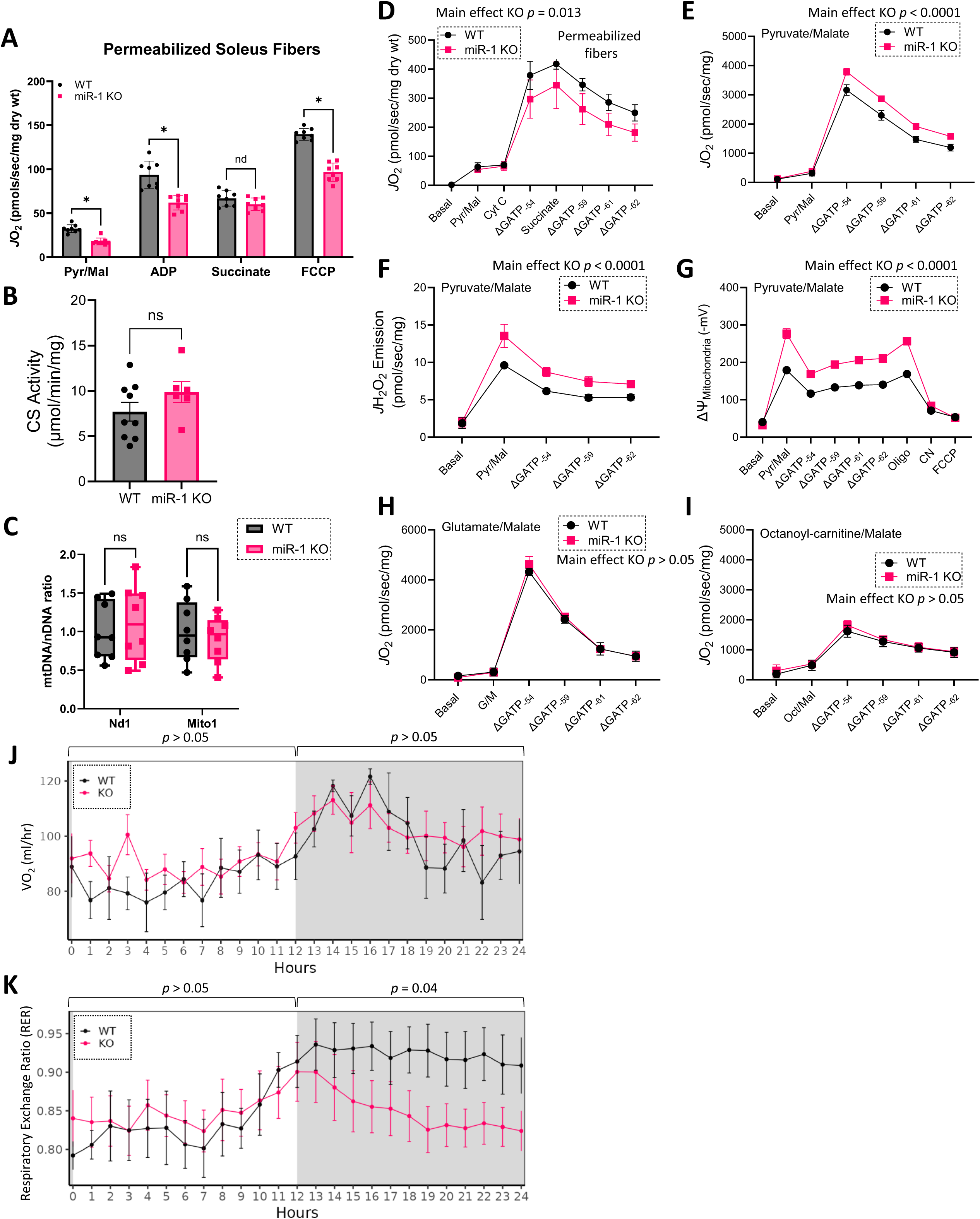
Bioenergetic phenotyping of miR-1 KO skeletal muscle. **(A)** Assessment of mitochondrial respiration in permeabilized soleus fibers (n=8 mice per group, post-tamoxifen treatment and an 8-week washout). Oxygen flux (*J*O2) normalized to mg tissue dry weight. Pyruvate/Malate-supported complex I leak (Pyr/Mal), ADP-stimulated OXPHOS (ADP), complex II OXPHOS (Succinate), and maximum respiration/*ET* capacity (FCCP) tested. **(B)** Citrate synthase (CS) activity in gastrocnemius complex lysates, normalized to protein **(C)** Quantitation of mitochondrial DNA (mtDNA) by levels of NADH Dehydrogenase 1 (Nd1), and Mito1 relative to nuclear DNA (nDNA) in gastrocnemius muscle. Data in **(B-C)** from n=6-9 female mice per group, post-tamoxifen treatment and an 8-week washout. **(D)** Assessment of OXPHOS kinetics using the CK clamp technique (increasing free energies [i.e., more negative ΔGATP values correspond to an increased ATP/ADP ratio)] with Pyr/Mal and succinate as substrates in permeabilized fibers from the gastrocnemius, data normalized to mg dry weight. **(E)** Assessment of OXPHOS kinetics using the CK clamp technique with Pyr/Mal as substrates in isolated mitochondria, data normalized to total protein. **(F)** H2O2 emission rate (*J*H2O2) assessed in isolated mitochondria in response to Pyr/Mal using a CK clamp, normalized to total protein. **(G)** Mitochondrial membrane potential (ΔΨ), expressed in mV, in response to Pyr/Mal using a CK clamp. Oligo: oligomycin, CN: cyanide. **(H)** OXPHOS kinetics using the CK clamp technique with glutamate/malate (G/M) as substrates in isolated mitochondria, data normalized to total protein. **(I)** OXPHOS kinetics using the CK clamp technique with octanoyl-carnitine/malate (Oct/Mal) as substrates in isolated mitochondria, data normalized to total protein. Data in **(D-I)** from n=4 WT and n=4 miR-1 KO female mice (post-tamoxifen treatment and an 8-week washout), and data are mean ± SEM. Data in **(A-C)** analyzed by *t*-tests or multiple *t*-tests, ns: not significant, \**p* < 0.05. Data in **(D-I)** analyzed by two-way ANOVA, main effect of miR-1 KO shown. **(J)** Whole-body oxygen consumption (VO2) and **(K)** Respiratory exchange ratio (RER) via indirect calorimetry. Data are mean ± SEM, n=5 female mice per group (post-tamoxifen treatment and an 8-week washout). Data analyzed using a two-way ANOVA for light and dark cycles separately.

The lower mitochondrial respiration of permeabilized miR-1 KO myofibers could be due to reduced mitochondrial content or metabolic suppression independent of mitochondrial content/density. Therefore, we measured citrate synthase (CS) activity and the ratio of mitochondrial DNA (mtDNA) to nuclear DNA (nDNA) in gastrocnemius complex (gastrocnemius, soleus, and plantaris) samples as a proxy for mitochondrial content. We found no difference between WT and miR-1 KO in either measure of mitochondrial content (Figures 3B-C). Thus, similar to a recent study demonstrating reduced pyruvate oxidation in sedentary mice without a reduction in mitochondrial density [63], our data further support the idea that lower muscle mitochondrial content is not necessary for the observed effects on glucose metabolism.

To further interrogate mitochondrial respiration under more physiological conditions, we applied a modified version of the creatine kinase (CK) energetic clamp to assess mitochondrial energy transduction across a range of energy demands [64–68]. In permeabilized gastrocnemius fibers, miR-1 KO mice displayed significantly reduced Pyr/Mal-stimulated mitochondrial respiration across a range of ATP free-energy states (23.41% lower on average, *p* = 0.013, Figure 3D). We next tested different respiratory substrates in isolated mitochondria prepared from the gastrocnemius complex. In contrast to the findings from permeabilized fibers, in isolated mitochondria, in which intracellular interactions are not preserved and glycolytic enzymes are not present [69], Pyr/Mal-stimulated respiration was significantly higher in miR-1 KO mitochondria compared to WT across increasing ATP free energies (26.66% higher on average, *p* < 0.0001, Figure 3E). Pyr/Mal-stimulated respiration was also associated with elevated hydrogen peroxide emission (*J*H2O2) and hyper-polarization of the mitochondrial membrane potential (ΔΨ) across a span of ATP free energy in miR-1 KO mitochondria (39.03% and 49.31% higher on average, respectively, *p* < 0.0001 for both, Figures 3F-G). There was no difference between WT and miR-1 KO mitochondrial oxygen consumption with glutamate/malate (Figure 3H) or fatty acid substrates (octanoyl-carnitine/malate) (Figure 3I).

Altered pyruvate-stimulated respiration in the absence of a change in fatty acid-stimulated respiration has been reported to be a feature of pyruvate oxidation resistance and metabolic inflexibility [63]. A key feature of metabolic inflexibility is an inability to increase the respiratory exchange ratio (RER), an indicator of substrate oxidation preference, during the dark cycle when mice are active and shift to carbohydrate oxidation [63]. To determine RER over a 24 hr period, miR-1 KO and WT mice underwent indirect calorimetry to measure whole-body oxygen consumption (VO2) and carbon dioxide production. Although VO2 was not influenced by the loss of miR-1 (Figure 3J), miR-1 KO mice displayed a significantly lower shift to carbohydrate oxidation during the dark cycle, indicating mitochondrial pyruvate resistance and a state of metabolic inflexibility (*p* = 0.04, Figure 3K).

We combined these bioenergetic assessments with mitochondrial-targeted nano-liquid chromatography mass spectrometry (nLC-MS/MS) proteomics for further evaluation of mitochondrial enrichment [66]. Proteomic data analysis showed that the mitochondrial enrichment factor (MEF), calculated as the proportion of all quantified proteins that could be identified as mitochondrial using the MitoCarta 3.0 database [70], was not different between WT and miR-1 KO mitochondrial preparations (Figure 4A). Moreover, the percentage of the mitochondrial proteome devoted to each OXPHOS complex was not different between WT and miR-1 KO mitochondrial preparations when normalized to total protein (Figure 4B).

**Figure 4.**
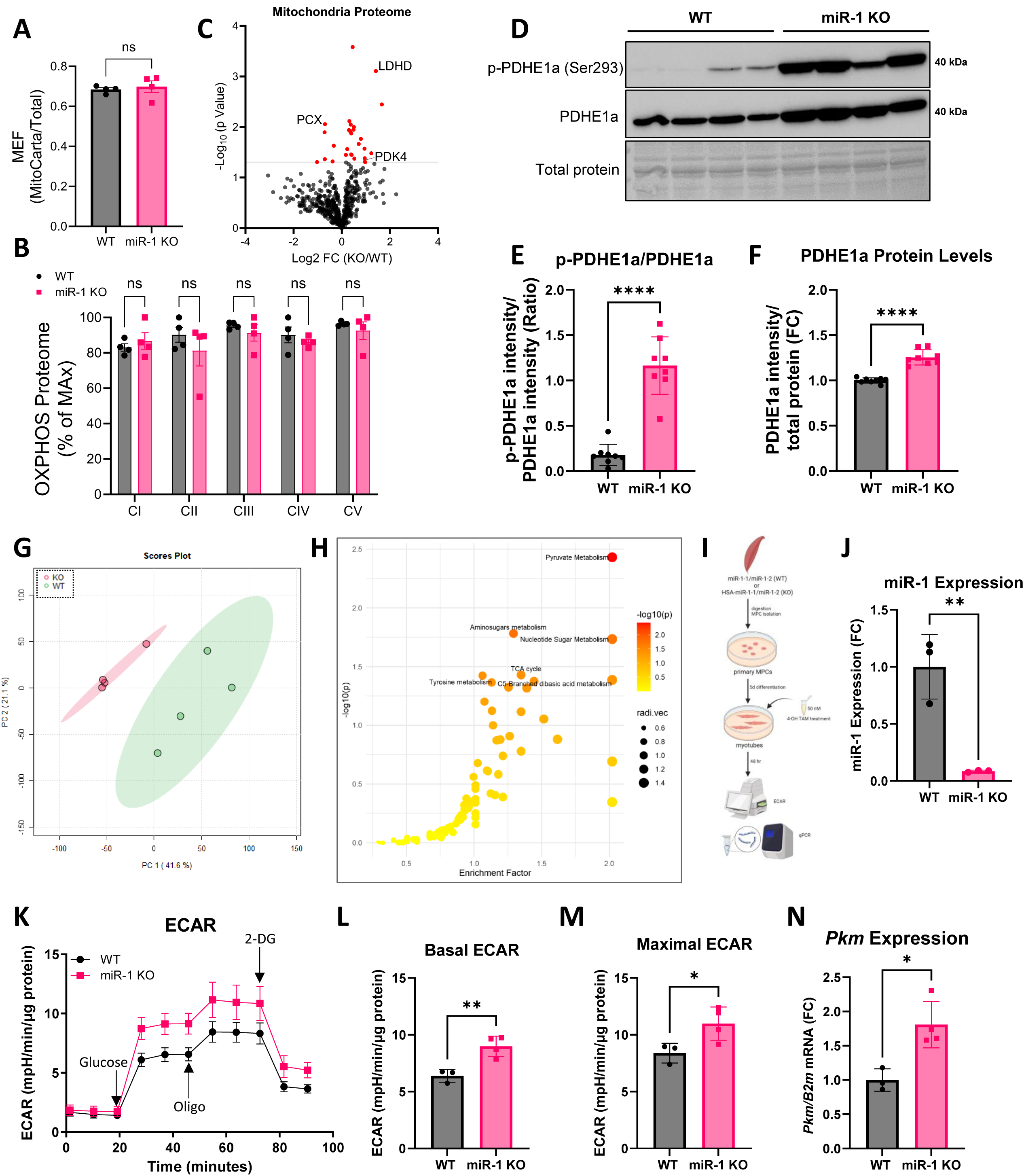
Mitochondrial proteomics and skeletal muscle metabolomic profiling of miR-1 KO skeletal muscle. **(A)** Ratio of mitochondrial protein to total protein abundance across samples, referred to as Mitochondrial Enrichment Factor (MEF). **(B)** Quantification of the OXPHOS protein complexes generated by the summed abundance of all subunits within a given complex. Data are presented as a percentage of the max for each complex. **(C)** Volcano plot depicting changes in the skeletal muscle mitochondrial proteome. Red color indicates significance (*p* < 0.05), and differentially expressed proteins involved in pyruvate metabolism are labeled. Data in **(A-C)** from n=4 WT and n=4 miR-1 KO female mice (post-tamoxifen treatment and an 8-week washout). **(D)** Representative Western blot of phosphorylation of Ser 293 on the PDHE1a subunit [p-PDHE1a(Ser293)], total PDHE1a, and corresponding total protein levels in WT and miR-1 KO gastrocnemius muscle lysates. **(E)** Quantification of p-PDHE1a/total PDHE1a protein levels after densitometric analysis of the levels of each sample normalized to corresponding total protein levels, expressed as a ratio. **(F)** Quantification of total PDHE1a protein levels after densitometric analysis of the levels of each sample normalized to corresponding total protein levels, expressed as FC. **(G)** Principal component analysis (PCA) scores plot generated using MetaboAnalyst based on gastrocnemius metabolites in n=4 WT and n=4 miR-1 KO female mice (post-tamoxifen treatment and an 8-week washout). Predictive component (PC) 1 and PC2 can differentiate the WT and miR-1 KO muscle. **(H)** Summary of altered metabolic pathways analysis with MetaboAnalyst reflecting the impact on the pathway and the level of significance. The colors of dots (varying from yellow to red) indicates the significance of the metabolites in the data, and the size of the dot is positively corelated with the impact of the metabolic pathway. Top 6 pathways labeled. **(I)** Study design for *in vitro* studies, created using Biorender.com. **(J)** miR-1 expression of 4-Hydroxytamoxifen (4-OH TAM)-treated myotubes from WT or miR-1 KO mice (n=3 untreated female mice per group). **(K)** Extracellular acidification rate (ECAR) trace over time after injection of indicated glycolytic modulators in WT (n=3 female) and miR-1 KO (n=4 female)-derived myotubes. Oligo: oligomycin, 2-DG: 2-deoxy-D-glucose. **(L)** Quantification of basal ECAR (before Oligo addition), and **(M)** maximal ECAR (after Oligo addition). **(N)** *Pkm* mRNA expression of WT and miR-1 KO myotubes. Data in **(E-F, J, L-N)** analyzed using independent *t*-tests. \**p* < 0.05, \*\**p* < 0.01, \*\*\*\**p* < 0.0001.

We next focused on pyruvate metabolism-related proteins that were differentially expressed in the mitochondrial-targeted proteomics. As shown in Figure 4C, lactate dehydrogenase D (LDHD) and pyruvate dehydrogenase kinase 4 (PDK4) were significantly up-regulated in miR-1 KO mitochondria compared to WT, while pyruvate carboxylase (PCX) was significantly down-regulated. To determine if higher PDK4 expression was associated with greater phosphorylation of pyruvate dehydrogenase (PDH), the protein levels of the E1α subunit of PDH that is phosphorylated at the Ser293 site (p-PDHE1a Ser 293) were assessed relative to total PDH levels (Figure 4D). Strikingly, we observed a significant up-regulation of the phosphorylation status of PDH (Figure 4E), as well as higher total PDHE1a levels (Figure 4F) in gastrocnemius muscles of miR-1 KO mice. The hyper-phosphorylation of PDH leads to enzyme inactivation which causes the conversion of pyruvate to lactate by preventing pyruvate from entering mitochondria [71]. This type of metabolic reprogramming is now recognized as causing a state of metabolic inflexibility [72–76]. To this end, we measured *ex vivo* muscle lactate secretion, which demonstrated a significant increase in buffer lactate levels in glycolytic extensor digitorum longus (EDL) muscles, but not in soleus muscles of miR-1 KO mice compared to WT (Supplemental Figures S4A-B). Taken together, these data suggest that high miR-1 levels are required for skeletal muscle to maintain metabolic flexibility.

### Loss of miR-1 alters the skeletal muscle metabolome

To determine if the change in pyruvate metabolism shown at the gene level in the miR-1 KO was observed at the metabolic level, we performed untargeted metabolomics on gastrocnemius muscle from WT (n=4) and miR-1 KO (n=4) mice. The metabolomic profiles between WT and miR-1 KO mice displayed clear and significant differences as observed by principal component analysis (PCA) (*p* = 0.024, Figure 4G). Pathway analysis of differentially expressed metabolites identified enrichment of pyruvate metabolism and TCA cycle metabolites, as well as metabolites involved in amino sugar and nucleotide sugar metabolism in miR-1 KO compared to WT (Figure 4H. These data provide compelling evidence to support that a primary function of miR-1 is the regulation of pyruvate metabolism in adult skeletal muscle and confirm our previous study proposing miR-1 regulation of the pentose phosphate pathway, as sugar backbones for all nucleotides are derived from ribose-5-phosphate in the pentose phosphate pathway [45].

Furthermore, integration of transcriptomic and metabolomic data of miR-1 KO mice demonstrated significant enrichment of glycolysis and pyruvate metabolism pathways (Supplemental Figure S4C). Notably, a number of glycolytic metabolites (glyceraldehyde 3-phosphate [GADP, Supplemental Figure S4D], 3-phosphoglycerate [3PG, Supplemental Figure S4E), phosphoenolpyruvate [PEP, Supplemental Figure S4F], lactic acid [Supplemental Figure S4G], and ribulose 5-phosphate [Ru5P, Supplemental Figure S4H]) were significantly up-regulated in miR-1 KO muscles.

### Loss of miR-1 leads to increased glycolysis in myotubes in vitro

To more closely examine glucose metabolism, we isolated myogenic progenitor cells (MPCs) from untreated WT or HSA-miR-1 mice at 6 months of age. Primary MPCs were allowed to differentiate for 5 days, and differentiated myotubes were then treated with 50 nM 4-Hydroxytamoxifen (4-OH TAM) *in vitro* to induce recombination in HSA-miR-1-derived myotubes (Figure 4I). Following 48 hours of treatment, myotubes were collected, and RNA was isolated to verify recombination and miR-1 KO (Figure 4J). Additionally, WT and miR-1 KO myotubes were analyzed using a Seahorse Bioanalyzer for extracellular acidification rate (ECAR) measurement, as a proxy for glycolysis (Figure 4K). Both basal and maximal ECAR were significantly higher in miR-1 KO myotubes compared to WT (Figures 4L-M). Moreover, qPCR analyses of primary myotubes identified significantly higher *Pkm* mRNA levels in miR-1 KO myotubes relative to WT (Figure 4N). Together, these data provide further support that loss of miR-1 increases glycolysis by targeting genes involved in the pyruvate metabolic pathway.

### Loss of miR-1 results in reduction of exercise performance conserved across species

Given the metabolic inflexibility observed in skeletal muscle of the miR-1 KO mouse, we hypothesized that exercise performance would be negatively impacted. To test this hypothesis, we first examined the exercise activity in *mir-1* mutant *C. elegans* which were reported to have reduced ATP levels in body-wall muscle [77]. Notably, the miR-1 seed sequence is highly conserved, and miR-1 is predominantly expressed in muscle tissues of *C. elegans* [78]. Adult *mir-1* mutant or N2 (WT) worms were transferred to an unseeded NGM plate (Untrained) or an unseeded NGM plate flooded with M9 buffer (Exercise) for two 90-minute sessions per day for 5 days, as previously described [79, 80] (Figure 5A). Locomotion analysis software [81, 82] used to record movement demonstrated that Untrained *mir-1* mutant worms had poorer swim performance parameters, including significantly lower wave initiation rate (Figure 5B) and significantly lower activity index compared to Untrained N2 worms (Figure 5C). Following the 5-day exercise protocol, wave initiation rate and activity index were significantly increased in N2 worms compared to Untrained N2 worms (Figures 5D-E). The *mir-1* mutant worms did not show a significant change in these physiological parameters in response to exercise compared to Untrained *mir-1* mutant worms. Additional physiological parameters measured are shown in Supplemental Figures S5A-F. To assess whether these functional effects are associated with altered pyruvate metabolism-related genes, we measured mRNA levels of *pdhk-2*, the *C. elegans* PDK ortholog. *Pdhk-2* expression was significantly higher in Untrained *mir-1* mutant worms compared to N2 worms (Figure 5F). However, the 5-day exercise protocol significantly reduced *Pdhk-2* expression in *mir-1* mutant worms.

**Figure 5.**
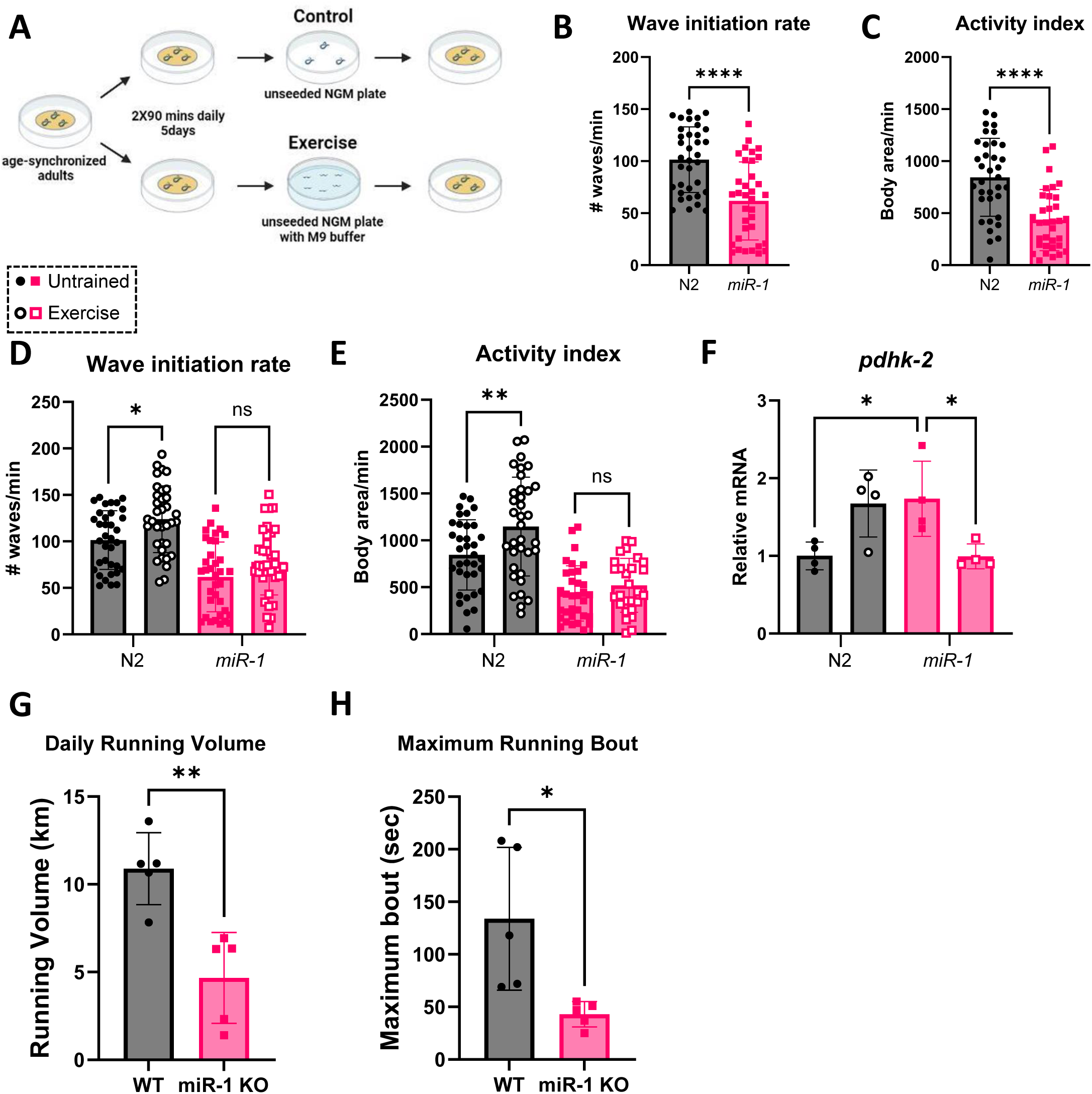
Effect of miR-1 loss on swimming activity in *C. elegans* and voluntary wheel running in mice. **(A)** Schematic of *C. elegans* swimming protocol. **(B-C)** CeLeST analysis of **(B)** wave initiation rate and **(C)** activity index in Untrained N2 (WT) (n=32) and *mir-1* mutant worms (n=35). **(D-E)** CeLeST analysis of **(D)** wave initiation rate and **(E)** activity index in Untrained (dark circle) or Exercised (open circle) N2 (n=32 and n=36, respectively), and Untrained (dark square) and Exercised (open square) *mir-1* mutant worms (n=35 and n=36, respectively). **(F)** *Pdhk-*2 mRNA expression in Untrained and Exercised N2 and *mir-1* mutant mice (n=4 per group), analyzed by qPCR. **(G-H)** ClockLab analyses of 4 weeks of voluntary wheel running. Average **(G)** daily running volume (km/day) and **(H)** maximum running bout (sec). Data in **(G-H)** from n=5 female mice per group (post-tamoxifen treatment and a 4-week washout). Data in **(B-C, G-H)** analyzed using independent *t*-tests, and data in **(D-F)** analyzed using multiple *t*-tests. Ns: not significant, \**p* < 0.05, \*\**p* < 0.01, \*\*\*\**p* < 0.0001.

We next investigated voluntary running wheel performance in adult miR-1 KO mice. At 4 months of age, mice were treated with tamoxifen, and following a 4-week washout, mice were singly housed with access to voluntary running wheels. After a week of acclimation, daily running volume and maximum running bout (longest continuous running period without stopping) were assessed over a period of 4 weeks. Consistent with the reduced exercise performance in the *mir-1* mutant *C. elegans*, we observed a significant reduction in running volume as well as maximum running bout in miR-1 KO mice compared to WT mice (Figures 5G-H). These data indicate that miR-1 has a deeply conserved role in facilitating endurance exercise performance.

### Down-regulation of miR-1 may play a role in rewired glucose metabolism during skeletal muscle hypertrophy in mice and humans

Ongoing research has provided evidence that in response to a hypertrophic stimulus, skeletal muscle undergoes a cancer-like metabolic reprogramming by shifting metabolism towards glycolysis [13, 83–87]. Previously, we reported that miR-1 expression is rapidly down-regulated by ∼70% in mice during mechanical overload (MOV)-induced skeletal muscle hypertrophy [88, 89]. We also found that during MOV-induced hypertrophy, pyruvate-stimulated mitochondrial respiration is significantly reduced [13]. Thus, while the present work demonstrates that the genetic and chronic loss of miR-1 in adult skeletal muscle results in metabolic inflexibility and reduced pyruvate-stimulated mitochondrial respiration, the physiological down-regulation of miR-1 in response to MOV may be a critical factor that allows muscle to undergo robust physiological hypertrophy. To explore this possibility, we compared genes up-regulated in miR-1 KO relative to WT, genes up-regulated in MOV relative to sham control [87], and our eCLIP-seq-identified miR-1 target genes. This integration identified 10 consensus miR-1 target genes that were up-regulated both during MOV and with miR-1 KO, which included *Ptbp1* (Figure 6A).

**Figure 6.**
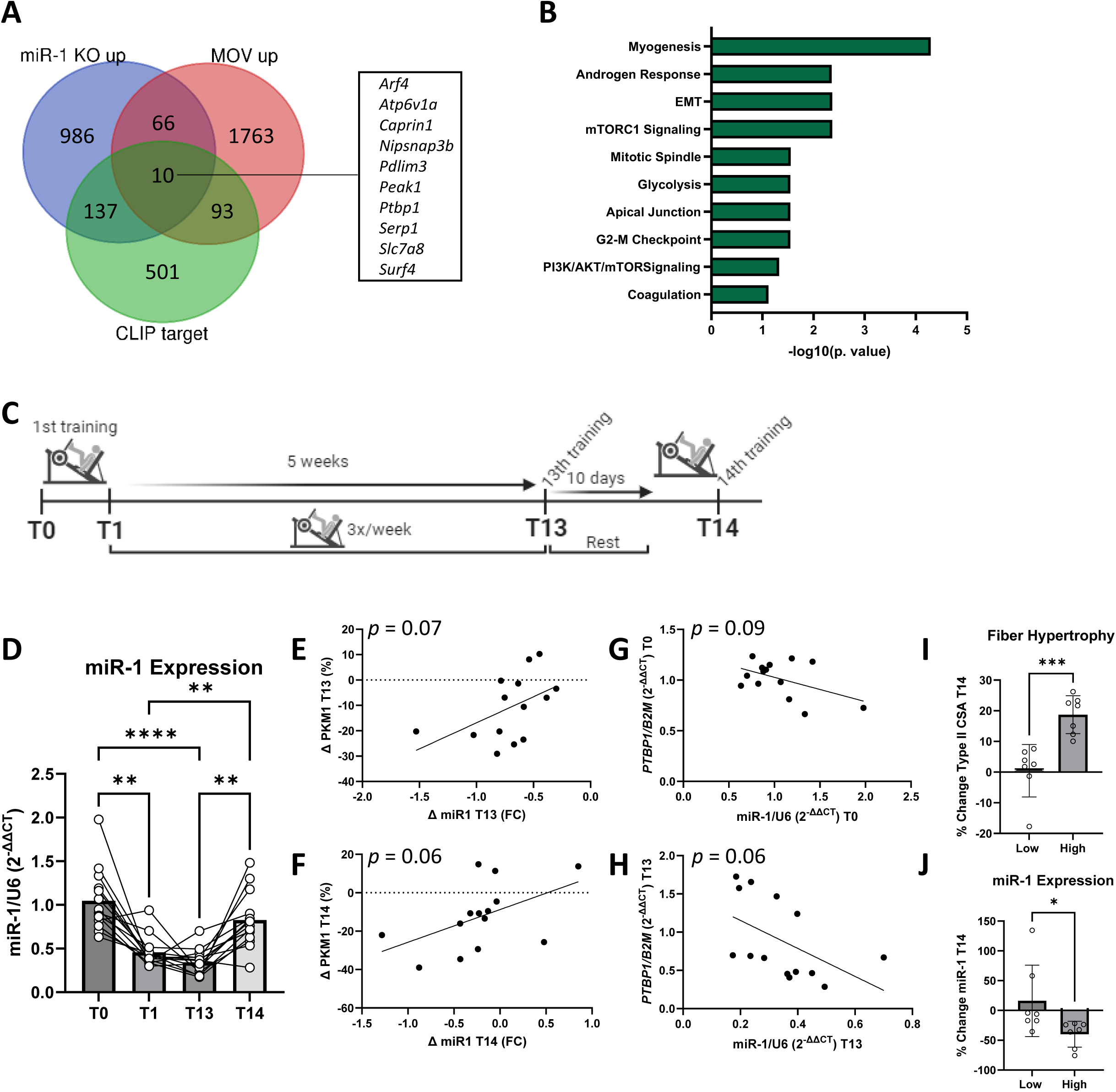
miR-1 down-regulation during MOV-induced muscle hypertrophy. **(A)** Venn diagram comparing significantly up-regulated genes in miR-1 KO compared to WT (“miR-1 KO up”), significantly up-regulated genes following 3 days of synergist ablation-induced MOV compared to sham (“MOV up”), and eCLIP-seq-defined miR-1 targets (CLIP target). Consensus genes listed. Venn diagram generated using https://bioinformatics.psb.ugent.be/webtools/Venn/. **(B)** Pathway enrichment analysis of consensus genes between “miR-1 KO up” and “MOV up,” top 10 pathways by adj. *p*-value shown. **(C)** Outline of human resistance exercise training program and times of biopsy collection (T0: baseline, T1: after the 1^st^ training session, T13: after the 13^th^ training session, T14: after the 14^th^ training session). **(D)** miR-1 expression in human skeletal muscle biopsies (from n=14 males) at the different time points of the resistance exercise training program. Differences in miR-1 expression at the different time points tested using a repeated-measures ANOVA with Tukey’s multiple comparisons. **(E-F)** Association between the change (Δ) in miR-1 expression and the Δ PKM1 protein levels at **(E)** T13 and **(F)** T14. **(G-H)** Association between miR-1 expression and *PTBP1* mRNA expression at **(G)** T0 and **(H)** T13. Associations tested using simple linear regressions, *p*-values shown. **(I)** Type II fiber hypertrophy, demonstrated as percent change in Type II fiber CSA from T0 to T14. Participants divided into Low (n=7) and High (n=7) groups based on magnitude of CSA increases. **(J)** Percent change in miR-1 expression from T0 to T14 in participants in the Low and High fiber hypertrophy groups. Data in **(I-J)** analyzed using independent *t*-tests Ns: not significant, \**p* < 0.05, \*\**p* < 0.01, \*\*\**p* < 0.0001.

Furthermore, pathway enrichment analysis of genes up-regulated in miR-1 KO and during MOV-induced hypertrophy found a common growth-related gene signature (i.e., mTORC1 Signaling, Glycolysis) (Figure 6B).

Similar to what we observed in the miR-1 KO, Verbrugge et al. previously demonstrated that six weeks of resistance exercise (RE) caused an increase in skeletal muscle PKM2 expression while PKM1 expression was down-regulated [90]. To determine if miR-1 expression was down-regulated in response to RE, we obtained human skeletal muscle biopsies from the participants of this prior study (Figure 6C) [90]. As shown in Figure 6D, miR-1 expression was significantly decreased after the first resistance training session (T1) compared to baseline (T0). The expression of miR-1 remained significantly lower after the 13^th^ training session (5 weeks, T13) but increased after 10-days of detraining. The reduction in miR-1 expression at T13 and T14 tended to correlate with the reduction in PKM1 at the corresponding time-point (*p* = 0.07 and *p* = 0.06, respectively, Figures 6E-F). In addition, higher miR-1 levels tended to correlate with lower levels of *PTBP1* both at baseline and at T13 (*p* = 0.09 and *p* = 0.06, respectively, Figures 6G-H). Finally, separating participants into “Low” (<10%, n=7) and “High” (≥10%, n=7) magnitude of muscle hypertrophy based on Type II fiber CSA changes from T0 to T14 (Figure 6I) demonstrated that the reduction in miR-1 was significantly greater in participants in the High fiber hypertrophy group (Figure 6J). These findings indicate that the miR-1>PTBP1>PKM2 axis is potentially conserved and may contribute to the metabolic reprogramming of skeletal muscle to a cancer-like metabolic state that promotes hypertrophic growth in response to mechanical loading.

## Discussion

Skeletal muscle-enriched miRNAs are increasingly being studied for their role in normal muscle function as well as in growth, atrophy, and disease [91, 92]. Specifically, myomiRs are attractive targets since they are so highly enriched in skeletal muscle and are not likely to have off-target effects when manipulated. Despite this, even the most recent miRNA studies in skeletal muscle [93, 94] rely on bioinformatic approaches to identify miRNA:target gene interactions, which are plagued by overestimating the number of miRNA binding sites and may predict interactions that are biologically irrelevant [95]. We therefore utilized AGO2 eCLIP-seq data to biochemically define the miRNA:target gene network in adult skeletal muscle. Since miR-1 is the most abundant miRNA in adult skeletal muscle, we focused our efforts on defining the miR-1 target gene network, identifying 1286 miR-1 binding sites located within 1136 genes. By combining the AGO2 eCLIP-seq miR-1 interactome and RNA-seq data from an inducible, skeletal muscle-specific miR-1 KO mouse, we provide compelling evidence that miR-1 has a central role in maintaining the metabolic flexibility of adult skeletal muscle through post-transcriptional regulation of metabolic genes involved in pyruvate oxidation. The loss of metabolic flexibility upon the genetic inactivation of miR-1 leads to diminished endurance exercise performance but conversely may promote a metabolic reprogramming of the muscle to a cancer-like metabolic state that supports hypertrophic growth.

Although several genes regulate the glycolytic pathway, PKM1/2 are the rate-limiting glycolytic enzymes [96]. The different isoforms of PKM have distinct roles; whereas PKM1 promotes pyruvate flux to the TCA cycle, PKM2 promotes glycolysis [97]. PKM2 can also be aggregated into a dimeric form, which induces higher nucleic acid synthesis through the pentose phosphate pathway [98]. PKM2 is expressed primarily in proliferating cells and tumor cells [99]. The different PKM isoforms are produced by alternative splicing, under the regulation of different splicing factors, including PTBP1. Specifically, PTBP1 acts as an exonic splicing silencer, binding to an optimal motif located at intron 8 in the PKM mRNA, blocking inclusion of exon 9, and resulting in PKM2 expression by inclusion of exon 10 [60, 100]. PTBP1 expression is promoted by oncogenic transcription factors, including MYC [60]. Surprisingly, studies found that miR-1 is down-regulated in diverse types of cancers including gastric, colorectal, breast, prostate, and lung cancer, which leads to elevated PTBP1 and PKM2 expression [19, 50–52, 101–103]. Conversely, over-expression of miR-1 has been shown to attenuate cancer cell growth [19, 102, 104–107]. Beyond its role in cancer, few studies have assessed the role of PKM2 in skeletal muscle. In myogenic progenitor cells (MPCs), PKM2 was shown to be important for proliferation *in vitro* [108]. In differentiated myotubes, loss of PKM2 results in reduced myotube size under basal and IGF-1-stimulated conditions [90]. PKM2 expression in myotubes also leads to reductions in oxidative metabolism [109]. In the current study, we found that miR-1 targets both *Pkm and Ptbp1* and that loss of miR-1 was sufficient to elevate PTBP1 levels, modulating *Pkm* alternative splicing and promoting the Warburg-related PKM2 isoform. In addition to the PTBP1/PKM2 mechanism, our AGO2 eCLIP-seq analyses revealed miR-1 binding sites that are responsive to genetic deletion of miR-1 involved in glycolysis and pyruvate metabolism, including *H6PD*, *PFKFB*, and *SLC16A3* (which encodes Monocarboxylate transporter 4, MCT4). H6PD, the endoplasmic reticulum (ER)-localized counterpart of glucose-6-phosphate dehydrogenase (G6PD), has been shown to play a critical role in pentose phosphate pathway activity of cancer cells [110]. PFKFB catalyzes the synthesis of fructose-2,6-bisphosphate (which acts as an allosteric regulator of phosphofructokinase [PFK]), and PFKFB has also been shown to regulate glycolysis in cancer cells [111, 112]. Lastly, MCT4 expression has been extensively characterized in cancer [113] and has also been shown to play a role in cardiac hypertrophy [114, 115].

Comprehensive bioenergetic phenotyping demonstrated pyruvate oxidation resistance with the loss of miR-1, which was further supported by evidence of PDH hyperphosphorylation. A number of studies have shown that inactivation of PDH results in a loss of metabolic flexibility in both cardiac and skeletal muscle [74, 76, 116–120]. The inability of miR-1 KO to switch to carbohydrate oxidation during their activity period, as reflected in a lower RER, provided additional evidence of metabolic inflexibility. Strikingly, the altered mitochondrial bioenergetics in miR-1 KO skeletal muscle was also accompanied by a significantly distinct metabolomic profile. Similar to the transcriptomic signature of the Warburg effect and metabolic reprogramming in cancer observed in miR-1 KO muscle, the metabolomic profile also demonstrated enrichment of pathways involved in cancer, including glycolysis and nucleotide sugar metabolism [121].

Metabolites up-regulated in miR-1 KO skeletal muscle included the glycolytic intermediates glyceraldehyde 3-phosphate, 3-phosphoglycerate, phosphoenolpyruvate, pyruvate, and lactate. Ribulose 5-phosphate (Ru5P), an end-product of the pentose phosphate pathway that is an essential component of purine and pyrimidine nucleotides biosynthesis [122], was also significantly up-regulated in miR-1 KO skeletal muscle. Figure 7 summarizes the effect of miR-1 KO on the glycolytic and pentose phosphate pathways, as demonstrated by RNA-seq and metabolomics. As shown, several glycolytic enzymes and metabolites were significantly up-regulated in skeletal muscle with the loss of miR-1. Moreover, a number of glycolytic genes were identified by AGO2 eCLIP-seq as miR-1 targets. Thus, we propose that the metabolic phenotype caused by the loss of miR-1 is the result of targeting multiple mRNAs of the molecular pathways of glycolysis and pyruvate metabolism rather than a single gene [123].

**Figure 7.**
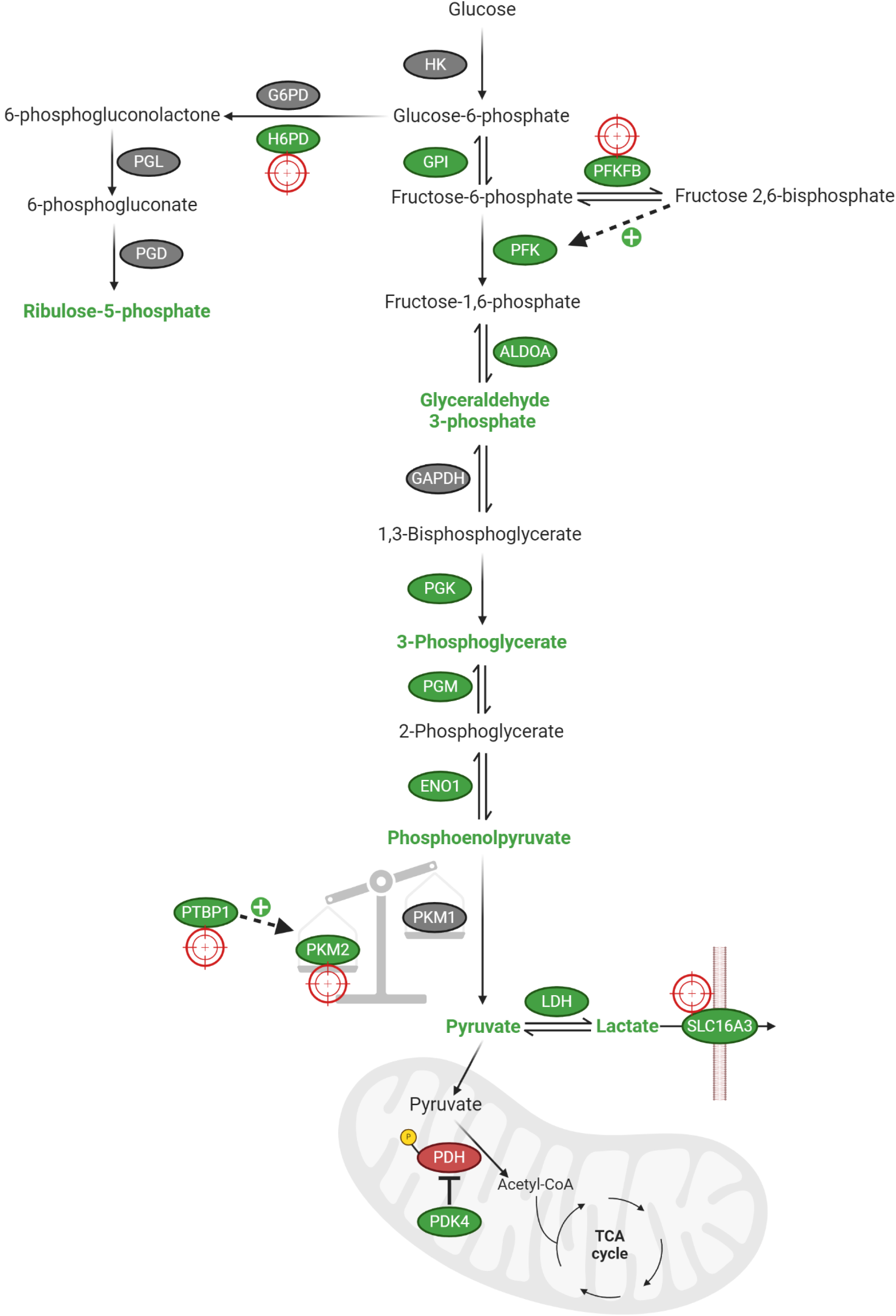
Summary of the effect of miR-1 loss on glycolytic and pentose phosphate enzymes and intermediates. Glycolytic and pentose phosphate pathway enzymes (ovals) and intermediates depicted. Enzymes up-regulated in miR-1 KO RNA-seq or mitochondrial proteomics indicated by green color, and inactivated enzymes indicated by red color. Up-regulated metabolites in miR-1 KO metabolomics indicated by bolded green text. Green plus sign indicates positive regulation. Target symbol indicates eCLIP-seq-defined miR-1 target. Created using Biorender.com.

Beyond transcriptomic, proteomic, and metabolic differences, the loss of miR-1 in skeletal muscle functionally resulted in reduced endurance exercise capacity in both *C. elegans* and in mice. miR-1 is highly enriched in cardiac and skeletal muscle, which are two organs that display unique metabolic adaptations to support their energy demands [35]; thus miR-1 may play a conserved role in regulating metabolic flexibility in these tissues. Interestingly, studies in cardiac muscle have also demonstrated metabolic and mitochondrial-related perturbations with miR-1 knockdown or KO [47, 124]. Although the loss of miR-1 has also been implicated in cardiac hypertrophy [47, 125–129], we found no difference in the muscle size of miR-1 KO compared to WT mice. Therefore, the metabolic reprogramming by miR-1 may not be sufficient to induce hypertrophy and may also require mechanical contraction. Recent studies increasingly suggest that hypertrophying skeletal muscle cells display a Warburg-like rewiring of metabolism [13, 83–86, 130]. Notably, miR-1 expression is dramatically down-regulated in response to a hypertrophic stimulus in mice [88, 89]. In the current study, we found that miR-1 is decreased during resistance exercise training-induced muscle hypertrophy in humans as well. Thus, while down-regulation of miR-1 may not be sufficient for muscle hypertrophy, a reduction in miR-1 and the ensuing metabolic reprogramming may be necessary for myofiber growth. Future research should investigate whether preventing the MOV-mediated down-regulation in miR-1 attenuates skeletal muscle hypertrophy. While it may appear counterintuitive that mitochondrial OXPHOS is downregulated during muscle growth, it is important to note that while mitochondria provide cellular energy by coupling substrate oxidation to ATP synthesis, this neglects other pathways of energy transduction, including NADPH, which can be reduced by the pentose phosphate pathway [131]. NADPH is also necessary as a reducing equivalent for nucleic acid and fatty acid synthesis. In addition, glycolysis is an efficient pathway, as most of the free energy of glucose metabolism to lactate is conserved in ADP phosphorylation to ATP [131]. Lastly, glycolysis produces less heat per ATP molecule than mitochondria [131]. Since maximally exercising skeletal muscle can be sensitive to heat [132], reducing heat production may be an additional benefit of switching to aerobic glycolysis during MOV-induced muscle hypertrophy.

Previous studies have provided evidence suggesting miR-1 is involved in regulating skeletal muscle metabolism [24, 36, 133]. While these studies proposed different mechanisms by which miR-1 regulates muscle metabolism, they were limited by the lack of bona fide miR-1 target genes identified by our AGO2 eCLIP-seq data. This limitation may have prevented the identification of our newly described pathways. Interestingly, intramuscular injection of miR-1 was shown to improve insulin resistance [133], which could have been due to the direct effect of higher pyruvate oxidation via the miR-1 mediated mechanisms described here. Furthermore, we found that extracellular vesicle delivery of miR-1 from skeletal muscle to adipose tissue, or transfection of miR-1 into adipocytes directly, promotes adrenergic signaling and lipolysis in adipocytes [89], which may involve effects on pyruvate metabolism [134]. Thus, the over-expression of miR-1 in non-muscle cells (i.e., cancer cells, adipocytes) may represent a novel approach to modulate the metabolism of these tissues.

It is important to note that one limitation of our multi-omics approaches was maintaining equal depth of coverage across the transcriptome for each methodology. While the advent of eCLIP-seq has dramatically reduced deduplication rates and input requirements from its predecessor, high-throughput sequencing of RNA isolated by crosslinking immunoprecipitation (HITS-CLIP), eCLIP in tissues still proves to be challenging. Therefore, we acknowledge that a percentage of differentially expressed genes identified by RNA-seq may possess miR-1 target-sites but were below our detection limits for peak calling. Additionally, our AGO2 eCLIP-seq peak-calling method uses neighboring genome read distribution to assess enrichment over background, which may mask weaker miR-1 bound states in favor of other non-miR-1 binding events.

Nonetheless, our studies identified, with high confidence, several miR-1 target genes involved in the glycolytic and pyruvate metabolic pathways. In this regard, miR-1 represents a single miRNA that can regulate entire biologic pathways by modulating several functionally related genes [135]. Finally, in addition to identifying a novel role for miR-1 in adult skeletal muscle tissue, our studies provide a foundation for future skeletal muscle miRNA research. AGO2 eCLIP-seq data, for example, as provided by the human skeletal muscle eCLIP-seq miRNA binding map, will guide future miRNA research to ensure the identification of biologically-relevant target genes.

## Methods

### AGO2 eCLIP and data processing

The AGO2 eCLIP (enhanced crosslinking and immunoprecipitation) protocol was executed with minor modifications to previously described methods [136, 137]. Skeletal muscle tissue was obtained from the University of Kentucky Center for Muscle Biology (CMB) biobank. Human samples were collected with written informed consent from each participant and the approval of the Institutional Review Board of the University of Kentucky. Participants’ written consent forms and the study protocol conformed to the ethical guidelines of the 1975 Declaration of Helsinki. Muscle biopsies were derived from the vastus lateralis of participants (n=3 males, n=3 females, 24.43 ± 5.35 y).

Skeletal muscle tissues were pulverized and subjected to UV-irradiation to crosslink protein-nucleic acid interactions. RNA associated with AGO2 was immunoprecipitated using a custom human anti-rabbit AGO2 antibody [138], followed by adapter ligation and RT-PCR to create libraries for high-throughput sequencing. Single-end 100 bp reads were performed on an Illumina Hi-Seq (Illumina, San Diego, CA, USA). Sequencing reads were pre-processed and mapped with minor changes to established protocols. Adapter sequences were trimmed, and the resulting reads were mapped to the human genome (hg19). Identical alignments were collapsed to eliminate PCR duplicates, and strand-specific read coverage was calculated from the alignments of the six samples.

To identify significant AGO2 clusters, we applied a zero-truncated negative binomial (ZTNB) model to calculate the significance of read coverage at each mapped genomic position [41]. The assumption was that read heights for each gene are sampled from an underlying ZTNB distribution. Parameters for gene-specific ZTNB probability density functions were estimated using read heights measured at all positions within an annotated gene region for each sample. *P*-values were calculated based on the probability of observing a read height as large as the observed height and were assigned to each position. Fisher’s method was used to summarize *p*-values at each genomic position across the six samples. Positions with a false discovery rate (FDR) of less than 5% were considered significant. Significant positions within 60 nucleotides of each other were merged into single contiguous intervals, and regions shorter than 50 nucleotides were extended symmetrically to 50 nucleotides to account for the AGO2 binding footprint. Intragenic cluster positions were then annotated according to their overlapping gene structures using Ensembl v75 annotations.

HOMER software was used to discover motifs enriched in AGO2 eCLIP cluster sequences. Additionally, motif enrichment was performed using a sliding window approach, as done previously [41]. Briefly, a single base offset sliding window of 7 nts was used to determine heptamer frequencies in cluster and background sequences. Enrichment data were calculated for all possible heptamers. Enrichment data for miR-1 seed was calculated individually for enrichment of 7M8 and 7A1 sequences. Enrichment significance was calculated by Fisher’s exact test, and *p*-values were transformed to Benjamini-Hochberg FDRs. TargetScan version 8.0 and miRcode miR-1 genome coordinate predictions were intersected with significantly enriched clusters using bedtools. AGO2 eCLIP-seq data are available on UCSC Genome Browser (https://genome.ucsc.edu/s/bdpeck/Ago2%20eCLIP).

### Generation of inducible miR-1 KO mouse

All animal procedures were conducted in accordance with institutional guidelines for the care and use of laboratory animals and approved by the Institutional Animal Care and Use Committee of the University of Kentucky. Mice were housed in a temperature- and humidity-controlled room and maintained on a 14:10-hr light-dark cycle with food and water *ad libitum*. To knockout miR-1 specifically in adult skeletal muscle, the skeletal muscle-specific inducible *Cre* mouse (HSA-MCM) [48] was crossed with a floxed *miR-1-1 and miR-1-2* (*miR-1-1^f/f^; miR-1-2^f/f^*) mouse [47] to generate the HSA-MCM; *miR-1-1^f/f^; miR-1-2^f/f^* mouse, designated HSA-miR-1.

Adult (4 months of age) female HSA-miR-1 mice were administered tamoxifen (2 mg/day) by intraperitoneal injection for five consecutive days to induce KO of miR-1. Tamoxifen-treated littermate *miR-1-1^f/f^; miR-1-2^f/f^* mice (HSA-MCM negative) pr vehicle-treated (15% ethanol in sunflower seed oil) HSA-miR-1 mice served as WT controls. Unless otherwise noted, after an 8 week-washout, (6 months of age), WT and KO mice were subjected to downstream experimentation or humanely euthanized via carbon dioxide asphyxiation followed by cervical dislocation. Hindlimb muscles were then carefully excised and prepared for storage or immediate downstream analyses, as described in sections below.

### Genotyping

For *miR-1-1^f/f^* genotyping, the following forward (F) and reverse (R) primers were used: F: 5’-GGG AGC GGT TCC TTA CGA CCA TC-3’; R: 5’-TCC ATC GGT CCA TTG CCT TTC-3’. DNA samples for genotyping were ran with a negative control and positive control. The PCR product for the *miR-1-1* WT allele was 1000 bp, while the *miR-1-1^f/f^* allele generated a larger PCR product (∼1100 bp). For *miR-1-2^f/f^*genotyping, genotyping primers were as follows: F: 5’-CGC AGG AGT GCC TAC TCA G-3’ and R: 5’-AGT GCT GAA GAT AGC ACT TGC C-3’. The PCR product for the *miR-1-2* WT allele was <1500 bp (∼1400 bp), while the *miR-1-2^f/f^* allele generated a larger PCR product 1500 bp in size.

### Gene expression

For RNA-seq and qPCR, total RNA was isolated from muscle previously snap-frozen in liquid nitrogen. Samples were homogenized in TRIzol using a THb Handheld Tissue Homogenizer and Hard Tissue Omni Tip Plastic Homogenizer Probes (Omni International, Kennesaw, GA, USA). RNA was isolated using a Direct-zol RNA Miniprep Kit (R2053, Zymo Research, Irvine, CA, USA) according to manufacturer’s instructions. For RNA-seq, RNA samples were shipped to Novogene Corporation Inc. (Sacramento, CA, USA) for library preparation (polyA, non-strand specific) and sequencing (20 million PE150 reads, NovaSeq, Illumina, San Diego, CA). RNA-seq data were analyzed in R (Ver. 4.4) using the DESeq2 package [139]. Pathway analyses were performed using the Enrichr web-based tool [140–142]. For RNA-seq and pathway analyses, significance was set using an FDR alpha of 0.05. For the CDF generated, miR-1 targets were tested against non-miR-1 targets using a Kolmogorov–Smirnov goodness-of-fit test. RNA-seq data have been deposited in the Gene Expression Omnibus (GEO) database (GSE274378). For comparison of miR-1 KO RNA-seq data with RNA-seq data during muscle hypertrophy, the list of differentially expressed genes from a previous study of 72 h of synergist ablation-induced MOV was downloaded [87]. Raw MOV data are available in GSE213406.

For gene expression of mRNAs, cDNA was synthesized using a High-Capacity RNA-to-cDNA kit (4387406, Thermo Fisher, Waltham, MA, USA). Prepared cDNA templates were analyzed by qPCR using TaqMan probes (4331182, Thermo Fisher, *Mib1*: Mm00523008_m1, *Pkm*: Mm00834102_gH; 4351372, Thermo Fisher, *PTBP1:* Hs00914687_g1) and TaqMan Fast Advanced Master Mix (4444556, Thermo Fisher) run on a QuantStudio 3 PCR system (Thermo Fisher). Threshold cycle (Ct) values between these genes and the housekeeping gene beta-2-microglobulin (*B2m,* 4331182, Thermo Fisher, Mm00437762_m1) or 18S rRNA (4331182, Thermo Fisher, Hs99999901_s1) was used to normalize data, and the 2^(-delta delta Ct) method was used to calculate fold change.

For miRNA expression, cDNA was synthesized using a TaqMan MicroRNA Reverse Transcription Kit (4366596, Thermo Fisher) and TaqMan MicroRNA RT primers (4427975, Thermo Fisher). Gene expression of miRNAs was analyzed by qPCR using TaqMan MicroRNA Assays (4427975, Thermo Fisher) as follows: miR-1 Assay ID 002222, miR-133a Assay ID 002246, miR-206 Assay ID 000510. The endogenous control U6 snRNA (Assay ID 001973) was used for normalization, and the 2^ (-delta delta Ct) method was used to calculate fold change.

### Single fiber isolation

Isolation of single fibers for MuSC removal was based on the paper from Rosenblatt et al. [143]. EDL muscles were carefully dissected and handled via the tendons. Muscles were immediately placed into warmed collagenase solution (2 mg/ml Type I collagenase [MB-122-0050, Rockland Immunochemicals, Limerick, PA, USA], 1% penicillin-streptomycin [Pen-Strep, 97063-708, VWR, Radnor, PA, USA] in DMEM [30-2002, ATCC, Manassas, VA, USA]), and incubated at 35°C for 45 min, inverting the tube every 5-10 min. To stop digestion, muscles were transferred using a large bore pipette coated with heat inactivated horse serum (26050088, Thermo Fisher) to 60 mm culture plates (430166, Corning, Corning, NY, USA) pre-coated with heat inactivated horse serum to prevent fibers from sticking to the plates, and then filled with DMEM. Muscles were triturated with glass Pasteur pipettes until 20-30 fibers were released, then transferred to a new plate and placed in a 5% CO2 incubator at 37°C. 10% Matrigel Matrix-(354234, Corning) coated 24-well plates (3524, Corning) were prepared, then isolated fibers were retrieved from the incubator, and individual fibers were removed from suspension with normal Pasteur pipettes and placed in the center of Matrigel-coated wells. Fibers were allowed to settle and attach for 3 min to the Matrigel, and then 500 μl of plating medium consisting of 10% horse serum and 10 ng/ml basic fibroblast growth factor (bFGF) (354060, Corning) in DMEM was slowly added to wells, care being taken not to agitate the droplet in which the fiber was plated. Three days after plating, MuSCs were stuck to the Matrigel, and fibers were removed from the Matrigel without disturbing the surrounding cells by means of a Pasteur pipette pulled to a fine tip in flame. After fiber removal, fibers were immediately placed in TRIzol for RNA isolation.

### Western blotting

For Western blotting, muscle samples were homogenized in RIPA lysis buffer (97063-270, VWR) with protease and phosphatase inhibitor cocktails (97063-010, VWR and 786-450, G-Biosciences, St, Louis, MO, USA) using a THb Handheld Tissue Homogenizer. Protein concentrations were determined using a Pierce BCA Protein Assay Kit (23252, Thermo Fisher). Samples were then prepared in Laemmli sample buffer (1610737, Bio-Rad Laboratories, Hercules, CA, USA), boiled, and 20 μg of proteins were separated by SDS-PAGE using 4-20% Criterion TGX Stain-Free Protein Gels (5678094, Bio-Rad). Total protein stains were activated by UV excitation using a ChemiDoc MP system (Bio-Rad), and then proteins were transferred onto LF PVDF membranes (1620263, Bio-Rad). Membranes were blocked in 5% nonfat dry milk (1706404, Bio-Rad) for 1 hr at room temperature, followed by primary antibody incubation overnight at 4°C. The primary antibodies and respective concentrations used were as follows: anti-PKM1 (1:1000, 7067, Cell Signaling, Danvers, MA) anti-PKM2 (1:1000, 4053, Cell Signaling), anti-phospho-PDHE1a (Ser293) (1:1000, NB110-93479, Novus Biologicals, Centennial, CO, USA) and anti PDHE1a (1:1000, NBP2-33922, Novus Biologicals). Following primary antibody incubation, membranes were washed in TBS-0.1% Tween-20 and then incubated in secondary antibodies (1:10,000, goat anti-rabbit IgG H+L secondary antibody HRP conjugate, 31460, Thermo Fisher) at room temperature for 1 hr. Blots were developed with enhanced chemiluminescence using Clarity Western ECL Substrate (1705061, Bio-Rad), and imaged with a ChemiDoc MP system. Blots were quantified using Image Lab Software (Bio-Rad). Restore Western Blot Stripping Buffer (21059, Thermo Fisher) was used to strip membranes that were reprobed with a second primary antibody.

### Immunohistochemistry

Muscle fiber size and fiber-type distribution were assessed using methodology previously described by our lab [144]. After harvesting, soleus or plantaris muscles were embedded in Tissue TEK Optimal Cutting Temperature (4583, Sakura Finetek, Torrance, CA, USA) and frozen using liquid nitrogen chilled isopentane. Unfixed 7 μm thick cryosections placed onto SuperFrost Plus slides (12-550-15, Thermo Fisher) were incubated in primary antibodies against myosin heavy chain (MHC) Types I (BA.D5), IIA (SC.71), IIB (BF.F3) (1:100, Developmental Studies Hybridoma Bank, Iowa City, IA, USA) as well as anti-laminin (1:100, L9393, Sigma-Aldrich, St. Louis, MO, USA) for 90 min at room temperature. In order to visualize MHC and laminin, fluorescence-conjugated secondary antibodies were applied to different mouse and rabbit immunoglobulin subtypes for 1 hr at room temperature (1:250, A-21242, A-21121, A-21426, A-11046, Thermo Fisher). Type IIX fibers were inferred from unstained fibers. Muscle sections were imaged using a Zeiss upright microscope (AxioImager M1, Zeiss, Oberkochen, Germany) at 20x magnification and analyzed by MyoVision software [145, 146].

### Bioenergetic phenotyping

For assessment of mitochondrial respiration in permeabilized fibers [147], muscles were removed, cut lengthwise into small samples, and immediately placed in ice-cold biopsy relaxing and preservation solution (BIOPS: 10 mM Ca-EGTA buffer, 0.1 μM free calcium, 20 mM imidazole, 20 mM taurine, 50 mM K-MES, 0.5 mM DTT, 6.56 mM MgCl2, 5.77 mM ATP [A9062, Sigma-Aldrich], 15 mM phosphocreatine [PCr, P1937, Sigma-Aldrich], pH 7.1). Fiber bundles were mechanically separated using forceps in a petri dish on ice, then transferred into BIOPS solution containing saponin (S2149, Sigma-Aldrich) at a final concentration of 50 μg/ml. Fiber bundles were agitated in saponin solution on ice for 30 min, transferred to mitochondrial respiration medium (5 mM ATP [A9062, Sigma-Aldrich], 105 mM K-MES, 30 mM KCl, 10 mM KH2PO4, 5 mM MgCl2, 1 mM EGTA, 2.5 g/L BSA) for 10 min, and then fiber bundles of ∼1-2 mg wet mass were added to Oroboros O2k Oxygraph (Oroboros Instruments, Innsbruck, Austria) chambers. All respiration experiments were carried out at 37°C in a 2 ml reaction volume. For permeabilized soleus fiber experiments, mitochondrial respiration medium was supplemented with 20 mM creatine monohydrate (C3630, Sigma-Aldrich), and substrates conditions tested were Pyr/Mal (10/2 mM, P2256/M1000, Sigma-Aldrich) to assess complex I leak, 4 mM ADP (A5285, Sigma-Aldrich) to initiate OXPHOS, succinate/rotenone (10/0.01 mM, S3674/ R8875, Sigma-Aldrich) to assess complex II OXPHOS, and 250 nM titrations of carbonyl cyanide p-trifluoromethoxyphenylhydrazone (FCCP, C2920, Sigma-Aldrich) up to 1.5 µM to assess maximum respiration/*ET* capacity).

For the remainder of the respirometry experiments, buffer was supplemented with 5 mM creatine monohydrate, 1 mM PCr, and 20 U/ml creatine kinase (CK) (10736988001, Sigma-Aldrich) to determine steady-state oxygen consumption rates (*J*O2) ranging from near resting up to ∼95% of maximal using a modified version of the CK energetic clamp technique [64–68]. In these assays, the free energy of ATP hydrolysis (ΔGATP) was calculated based on known amounts of creatine, PCr, and ATP in combination with excess CK and the equilibrium constant for the CK reaction [64]. Substrate conditions tested were as follows: Pyr/Mal (5/2.5 mM), succinate/rotenone (10/0.005 mM), glutamate (G1501, Sigma-Aldrich)/Mal (10/2.5 mM), and octanoyl-carnitine (50892, Sigma-Aldrich)/Mal (0.2/2.5 mM). Following substrate addition, sequential additions of PCr to 6, 15, and 21 mM were performed to gradually slow *J*O2 back toward baseline.

For functional assays involving isolated mitochondria, differential centrifugation was employed to prepare the mitochondria. Gastrocnemius complexes (gastrocnemius, soleus, and plantaris) were excised and immediately placed in ice-cold Buffer A (10 mM EDTA in PBS, pH 7.4).

Tissue was minced and then incubated on ice for 5 min in Buffer A, supplemented with 0.025% trypsin. Skeletal muscle suspensions were then centrifuged at 200 x g for 5 min at 4°C to remove trypsin. Tissue pellets were then resuspended in Buffer B (50 mM MOPS, 100 mM KCl, 1 mM EGTA, 5 mM MgSO4), supplemented with 2 g/L BSA and homogenized with a Teflon pestle and borosilicate glass vessel. Homogenates were then centrifuged at 500 x g for 10 min at 4°C, and supernatant from each tissue was filtered through thin layers of gauze and subjected to an additional centrifugation at 10,000 x g for 10 min at 4°C. Mitochondrial pellets were washed in 1.4 ml of Buffer B, then transferred to microcentrifuge tubes and centrifuged at 10,000 x g for 10 minutes at 4°C. Buffer B was aspirated from each tube, and the final mitochondrial pellets were suspended in 100-200 µl of Buffer B. Yield was assessed by protein concentration using the BCA method. Cytochrome c (10 μM) was included in all assays to check the integrity of the outer mitochondrial membrane.

Fluorescent determination of ΔΨ was carried out using a QuantaMaster Spectrofluorometer (QM-400, Horiba Scientific, Kyoto, Japan) via tetramethyl rhodamine-methylester (TMRM, T668, Thermo-Fisher), at 37°C as previously described [64]. The fluorescence ratio of the following excitation/emission parameters: Ex/Em (576/590)/(552/590) was recorded, and a KCl standard curve was used to convert the ratios to millivolts. Buffer C was supplemented with 10 μM cytochrome C and 0.2 μM TMRM, and isolated mitochondria (0.05 mg/ml) were added to the assay buffer. Respiratory substrates, CK clamp components were then added, followed by sequential PCr additions as in the respirometry experiments. Following the final PCr addition, 15 nM oligomycin (75351, Sigma-Aldrich) was added for hyper-polarization of the ΔΨ, and finally, 10 mM cyanide (60178, Sigma-Aldrich) was added to depolarize the ΔΨ.

Mitochondrial H2O2 emission (*J*H2O2) was measured fluorometrically via the Amplex Ultra Red (AUR)/horseradish peroxidase (HRP) system, as previously described [64, 148]. Fluorescence was monitored (Ex:Em 565:600) using a QuantaMaster Spectrofluorometer, and resorufin fluorescence was converted to pmoles H2O2 via an H2O2 standard curve. Assay buffer was buffer C, supplemented with 2.5 g/L BSA, 5 mM creatine, 1 mM PCr, 20 U/ml CK, 10 μM AUR (A36006, Thermo-Fisher), 1 U/ml HRP (P8375, Sigma-Aldrich), and 20 U/ml superoxide dismutase (S9697, Sigma-Aldrich). Isolated mitochondria (0.1 mg/ml) were added to assay buffer, followed by addition of respiratory substrates (Pyr/Mal), 0.1 μM auranofin (A6733, Sigma-Aldrich), 5 mM ATP, and sequential PCr additions of 6, 15, and 21 mM. For CK clamp experiments, differences between WT and miR-1 KO were tested using a two-way ANOVA for the main effect of genotype.

### Mitochondrial content

For measurement of CS activity in muscle lysates, a colorimetric plate-based assay was used, whereby CoA-SH, a byproduct formed by the CS-mediated reaction of oxaloacetate (OAA) and acetyl-CoA, interacts with 5′, 5′-dithiobis 2-nitrobenzoic acid (DTNB) to form TNB (OD: 412 nm). A 96-well round bottom plate was loaded with assay buffer C (105 mM K-MES, 30 mM KCl, 10 mM KH2PO4, 5 mM MgCl2, and 1 mM EGTA, pH 7.2) supplemented with 0.2 mM DTNB and 0.5 mM acetyl-CoA, and tissue lysates (10 µg/well), and then incubated at 37°C for 5 min to deplete endogenous substrates. The assay was then initiated by the addition of 1 mM OAA to sample wells, and absorbance was measured at 412 nm every 30 s for 20 min.

For mitochondrial DNA (mtDNA) analysis, total DNA was isolated from gastrocnemius muscles using a QiAmp DNA Mini kit (51304, Qiagen, Hilden, Germany), and 10 µg per reaction was used as a template for SYBR Green (Bio-Rad)-based qPCR, as previously described [149].

Mitochondrial target genes were NADH Dehydrogenase 1 (Nd1) (F: GGCTATATACAACTACGCAAAGGC, R: GGTAGATGTGGCGGGTTTTAGG) and Mito1 (F: ACATAGCACATTACAGTCAAATCCCTTCTCGTCCC, R: TGAGATTGTTTGGGCTACTGCTCGCAGTGC). H19 (F: CACTGGCCTCCAGAGCCCGT, R: CGTCTTGGCCTTCGGCAGCTG) was used as nuclear DNA control.

### Mitochondrial-targeted proteomics

Bioenergetic phenotyping was combined with subcellular mitochondrial-targeted nLC-MS/MS [66, 67, 150]. Isolated mitochondria were lysed in buffer consisting of 8 M urea in 40 mM Tris, 30 mM NaCl, 1 mM CaCl2, 1 x cOmplete ULTRA mini EDTA-free protease inhibitor tablet (05892953001, Roche Holding AG, Basel, Switzerland), pH 8.0. Samples were subjected to three freeze-thaw cycles, then sonicated with a probe sonicator in three 5 sec bursts at an amplitude of 30 (Q Sonica CL188, Qsonica, Newton, CT, USA). Samples were then centrifuged at 10,000 x g for 10 min at 4°C. Equal amounts of protein (determined by BCA protein assay) were reduced with 5 mM DTT at 37°C for 30 min, alkylated with 15 mM iodoacetamide at room temperature for 30 min in the dark, and unreacted iodoacetamide was quenched with DTT up to 15 mM. Digestion was initially performed with Lys-C Protease (90307, Thermo-Fisher, 1:100 w:w, 2 µg enzyme per 200 µg protein) for 4 hr at 37°C. Samples were diluted to 1.5 M urea with buffer containing 40 mM Tris (pH 8.0), 30 mM NaCl, 1 mM CaCl2, then digested overnight with 50:1 w/w protein:enzyme of trypsin (V5113, Promega, Madison, WI, USA) at 37°C. Samples were subsequently acidified to 0.5% TFA and centrifuged at 4000 x g for 10 min at 4°C. Supernatant containing soluble peptides was desalted, and the eluate was frozen and lyophilized.

Final peptides were resuspended in 0.1% formic acid, then quantified using a Pierce Quantitative Colorimetric Peptide Assay (23275, Thermo Fisher). Samples were diluted to a final concentration of 0.25 µg/µl, then subjected to nLC-MS/MS analysis using an Ultimate 3000 RSLCnano system (Thermo Fisher) coupled to a Q Exactive Plus Hybrid Quadrupole-Orbitrap mass spectrometer (Thermo Fisher) via a nanoelectrospray ionization source. For each injection, 1 µg of sample was initially trapped on a trapping column (Acclaim Pep Map 100, 200 mm x 0.075 mm, 164535, Thermo Fisher) 5 µl/min at 98/2 v/v water/acetonitrile with 0.1% formic acid. Analytical separation was then performed at a flow rate of 250 nl/min over a 95 min gradient of 4-25% acetonitrile using a 2 µm EASY-Spray PepMap RSLC C18 75 µm x 250 mm column (ES802A, Thermo Fisher) with a column temperature of 45°C. MS1 was performed at 70,000 resolution, with an AGC target of 3×10^6^ ions and a maximum injection time of 100 ms. MS2 spectra were collected by data-dependent acquisition of the top 15 most abundant precursor ions with a change greater than 1 per MS1 scan and dynamic exclusion enabled for 20 sec. MS2 scans were performed at 17,500 resolution, with an AGC target of 1×10^5^ ions and a maximum injection time of 60 ms. Precursor ions isolation window was 1.5 m/z, and normalized collision energy was 27.

Proteome Discoverer 2.2 (Thermo Fisher) was used for data analysis, with default search parameters, including oxidation and carbamidomethyl (15.995 Da on M and 57.021 Da on C, respectively) as variable and fixed modifications. Data were searched against the Uniprot Mus musculus reference proteome (UP 000000589) and the mouse Mito Carta 3.0 database [70]. Peptide spectrum matches (PSMs) were filtered to a 1% FDR and grouped to unique peptides while maintaining a 1% FDR at the peptide level. Peptides were grouped to proteins using rules of strict parsimony, proteins were filtered to 1% FDR. Peptide quantification was done using the MS1 precursor intensity, and imputation was performed via low abundance resampling.

Mitochondrial enrichment factor (MEF) was determined using only high confidence master proteins by comparing mitochondrial protein abundance (proteins identified as mitochondrial by cross-reference with the MitoCarta 3.0 database) to total protein abundance. Quantification of OXPHOS protein complexes was generated by the summed abundance of all subunits within a complex.

All mass spectrometry samples were normalized to total protein abundance, which was converted to the Log2 space. For pairwise comparisons between WT and miR-1 KO, tissue mean, standard deviation, and *p*-value (two-tailed Student’s *t*-test, assuming equal variance) were calculated.

### Indirect calorimetry

A Sable Promethion Core phenotyping system (Sable Systems International, North Las Vegas, NV, USA) was used for indirect calorimetry at 3-minute intervals. Metabolic data quantified included O2, CO2, and H2O vapor; energy expenditure was then calculated from the Weir equation, and the respiratory quotient (RQ) and RER was calculated as the ratio of the volume of CO2 produced to the volume of oxygen O2 used, or VCO2/VO2. The web-based tool CalR was used for indirect calorimetry analysis [151]. Mice were individually caged and acclimated for 24 hrs before data collection. Data were analyzed using a two-way ANOVA for light and dark cycles separately.

### Metabolomics

Frozen gastrocnemius samples, each weighing 25 mg were cryo-fractured in muscle sample extraction solution (50% HPLC-grade methanol and 100% formic acid in a 100:1 ratio) in tissueTUBE TT1 Extra Thick tubes (520007, Covaris, Woburn, MA, USA) using a Covaris CryoPREP CP02 Cryogenic Dry Pulverization system (Covaris). The pulverized samples were further homogenized with zirconium silicate beads (Next Advance, Inc., Troy, NY, USA) using a Bullet Blender Gold (Next Advance, Inc.). Samples were centrifuged at 20,000 x g for 10 min, then filtered using Captiva EMR-Lipid cartridges (Agilent Technologies). Filtered samples were collected in LC/MS autosampler vials (Agilent Technologies) by centrifugation at 500 x g for 20 min, followed by drying and concentration using a Savant SpeedVac Integrated Vacuum Concentrator System and Kit (SPD2030, Thermo Fisher). The sample pellets were resuspended in 5% acetonitrile, 0.1% formic acid, and 100% HPLC-grade water and then centrifuged at 6,000 x g for 30 sec. Samples were subsequently transferred to a centrifuge tube filter with a 0.22 µm pore (Costar, Glendale, AZ, USA) and centrifuged at 20,000 x g for 10 min at 4°C. The filtered samples were transferred into polymer feet autosampler inserts (Agilent Technologies) in LC/MS autosampler vials and stored at -20°C until LC-MS analysis.

For untargeted metabolomics, metabolites were separated using SeQuant ZIC-pHILIC columns (150 mm x 2.1 mm; 5 µm particle size, 150454, Sigma-Aldrich). Mobile phase A consisted of 10 mM ammonium acetate in water, pH 9.8, and mobile phase B consisted of 100% methanol. The column flow rate was set to 0.15 ml/min, and column temperature was set to 25°C. Metabolites were separated over a 19-minute gradient from 90% B to 30% B; the column was then washed for 5 min at 30% phase B, followed by re-equilibration at 90% B for 8 min. Metabolites were analyzed on an Orbitrap Exploris 240 mass spectrometer coupled to a Vanquish Neo UHPLC system (Thermo Fisher). For each polarity set, polarity-switching MS1 only acquisition was acquired at 120,000 FWHM resolution with a scan range of 80-1200 Da, an automatic gain control (AGC) target set to ‘Custom’ at 1E6 absolute AGC value, and maximum inject time of 50 ms. Compound Discoverer 3.3 SP (Thermo Fisher) was used for data analysis using the Untargeted Metabolite Processing Workflow. Data were searched against the following ChemSpider databases: BioCyc, Human Metabolome, and KEGG, to identify compound names. Metabolomic output was analyzed and visualized using the MetaboAnalystR R package [152].

To test significance of clusters on the Principal Component Analysis (PCA), a permutational multivariate ANOVA (PERMANOVA) was used. For differential expression of metabolites, *t*-tests were used. The MetaboAnalyst 6.0 web-based platform [153] was also used for joint pathway analysis using the gene list of differentially expressed genes from the RNA-seq together with the differentially expressed metabolite list [154]. For all metabolomics statistical analyses, FDR < 0.05 was used as the threshold for significance.

### Seahorse XF analysis

Hindlimb muscles from 6-month-old untreated *miR-1-1^f/f^; miR-1-2^f/f^* mice (WT) or HSA-miR-1 mice (miR-1 KO) were digested using a GentleMACS Octo Dissociator (130-096-427, Miltenyi Biotec, North Rhine-Westphalia, Germany) with a Skeletal Muscle Dissociation Kit, mouse and rat (130-098-305, Miltenyi Biotec), according to manufacturer’s instructions. Following digestion, MPCs were isolated using an autoMACS Pro Separator (130-092-545, Miltenyi Biotec) with a Satellite Cell Isolation Kit, mouse (130-104-268, Miltenyi Biotec). Primary MPCs were expanded on 10% Matrigel-coated (Corning) Primaria culture plates (353846, Corning) in growth media consisting of Hams F-10 (10-070-CV, Corning), 20% FBS (35-070-CV, Corning), 1% Pen-Strep (VWR), and 10 ng/ml bFGF (Corning). MPCs were split by mild trypsinization (L0154-01000, VWR) at around 40% confluence until time of differentiation. For differentiation to myotubes, MPCs were allowed to reach 85-95% confluency before switching from growth medium to differentiation medium, consisting of DMEM (30-2006, ATCC) supplemented with 2% horse serum (35-030-CV, Corning). MPCs were allowed to differentiate for 5 days before adding differentiation medium supplemented with 50 nM 4-OH TAM (89152-604, VWR) for 48 hrs to induce recombination in HSA-miR-1-derived myotubes (or for TAM control in WT-derived myotubes). Following 48 hrs of 4-OH TAM treatment, myotubes were subjected to either ECAR assessment or RNA isolation and qPCR as described above.

For ECAR experiments, culture and differentiation of myotubes were performed in Matrigel-coated Seahorse XF24 cell culture microplates (100882-004, Agilent Technologies, Santa Clara, CA). Before ECAR analysis using a Seahorse XF24 Analyzer (Agilent Technologies), media was replaced with Seahorse XF DMEM assay media (103575-100, Agilent Technologies). ECAR was assessed using a Seahorse XF Glycolysis Stress Test Kit (103020-100, Agilent Technologies). Immediately after ECAR measurement, protein concentrations were determined using a BCA assay, for protein normalization. All data were generated from four technical replicates (averaged) for each biological replicate (n=3 WT and n=4 KO).

### C. elegans swim exercise

*C. elegans* were cultured on NGM plates seeded with *E. Coli* (OP50) at 20°C. N2 (Bristol strain, wild type) and *mir-1* mutant MT17810 [*mir-1(n4102*) I] strains were obtained from Caenorhabditis Genetics Center (CGC), supported by the National Institutes of Health - Office of Research Infrastructure Programs (P40 OD010440). Swimming exercise was performed according to previous descriptions [79, 80]. Synchronized worms were bleached by sodium hydroxide bleaching buffer, then incubated overnight in M9 buffer on a rocker at 20°C. Larval stage L1 worm populations were transferred to 60 mm NGM plates seeded with OP50 for 2 days to obtain adult D1 worms, which were then washed off plates with 3 ml M9 buffer and allowed to settle under gravity, with the supernatant including OP50 and larvae removed, for a total of 3 times. Worms were then transferred to unseeded NGM plates with M9 buffer using a glass Pasteur pipette, and all plates were then moved to a 20°C incubator for 90 min. After swimming exercise, worms were washed off with M9 buffer, gravity settled, and transferred to new 60 mm NGM plates seeded with OP50 in a 20°C incubator. Swimming exercise was performed 2 x 90 min daily for 5 days.

CeleST (*C. elegans* Swim Test) was conducted as previously described [80–82]. One day following the 5-day swimming exercise regimen, a 10 mm pre-printed ring on the surface of a glass microscope slide was used and covered with 50 µl M9 buffer. 7-8 worms per group and 30 worms total were picked up and placed into the swimming area, and 60-sec movies with ∼16 frames per sec of the worms were taken using a Nikon LV-TC microscope (Nikon Instruments, Melville, NY, USA) at 1x magnification with an OPTIKA C-P20CM camera (Optika Microscopes, Ponteranica, Italy).

RNA isolation from *C. elegans* utilized TRIzol (Thermo Fisher) [Ref: 10.1101/pdb.prot101683]. Subsequent to RNA isolation, 500 ng of RNA was mixed with 1 μl of random hexamers (Sigma-Aldrich) and incubated at 65 °C for 10 minutes. The mixture was combined with a master mix including 4 μl of RT buffer (Sigma-Aldrich), 2 μl of DTT (Sigma-Aldrich), 1 μl of dNTP (Thermo Fisher), 1 μl of Superscript II (Sigma-Aldrich) and 1 μl of Ribolock (Thermo Fisher), followed by incubation at 42°C for 60 minutes. qPCR analysis was conducted using the SYBR Green Master Mix (Qiagen) in a 10 µl reaction volume. The quantification of gene expression, relative to the housekeeping gene CDC42, was determined utilizing the delta Ct method.

### Voluntary wheel running

Voluntary wheel cages for mice were from Actimetrics (Wilmette, IL, USA). Cage wheels (model PT2-MCR1) are 11-cm inside diameter, 5.4 cm wide, with 1.2-mm wide bars placed 7.5 mm apart. A wireless node was used to sense wheel revolutions from the rotation of a magnet mounted on the wheel axle, recorded via the ClockLab software (Actimetrics). For wheel running experiments, mice underwent a 4-week washout after tamoxifen treatment and then were single-housed in voluntary wheel cages for an additional 5 weeks. ClockLab analyses included number of wheel revolutions/day and bout length, as previously described [155]. The first week was used for acclimation, then data over the subsequent 4 weeks were averaged.

Daily running volume was defined as the average distance covered per day, and maximum bout was defined as the longest continuous running period without stopping.

### Human resistance exercise training samples

Human resistance exercise training vastus lateralis biopsy samples were collected in a previous study investigating skeletal muscle responses over the time course of 6 weeks of resistance exercise [156]. The lower limb exercise training regimen is described in detail in the original publication [156]. Briefly, 14 young males (24 ± 3 y) conducted thrice weekly resistance exercise for a total of 14 resistance exercise training sessions. Muscle biopsies were obtained at baseline (T0), after the first (T1), 13^th^ (T13), and after 10 days of rest, the 14^th^ session (T14).

The same samples were also used in another previously published study, which demonstrated that the protein abundance of PKM1 and PKM2 change in response to resistance exercise [90]. In the current study, we aimed to assess miR-1 levels in the muscle biopsy samples at the different time points. Biopsy samples were shipped on dry ice, and RNA isolation, miR cDNA synthesis, and qPCR were performed as described above.

### Statistical analyses

Unless otherwise specified above, Student’s *t* tests (2-tailed) were used to determine the significance between WT and KO groups. For human resistance exercise training samples, a repeated measures one-way ANOVA with Tukey’s multiple comparisons test was used. For correlation analyses, a simple linear regression was used. Unless otherwise stated, statistical analyses were performed using GraphPad Prism software (Ver. 10.2.3), and the level of significance was set at *p* < 0.05.

---

**Supplemental Figure S1.**
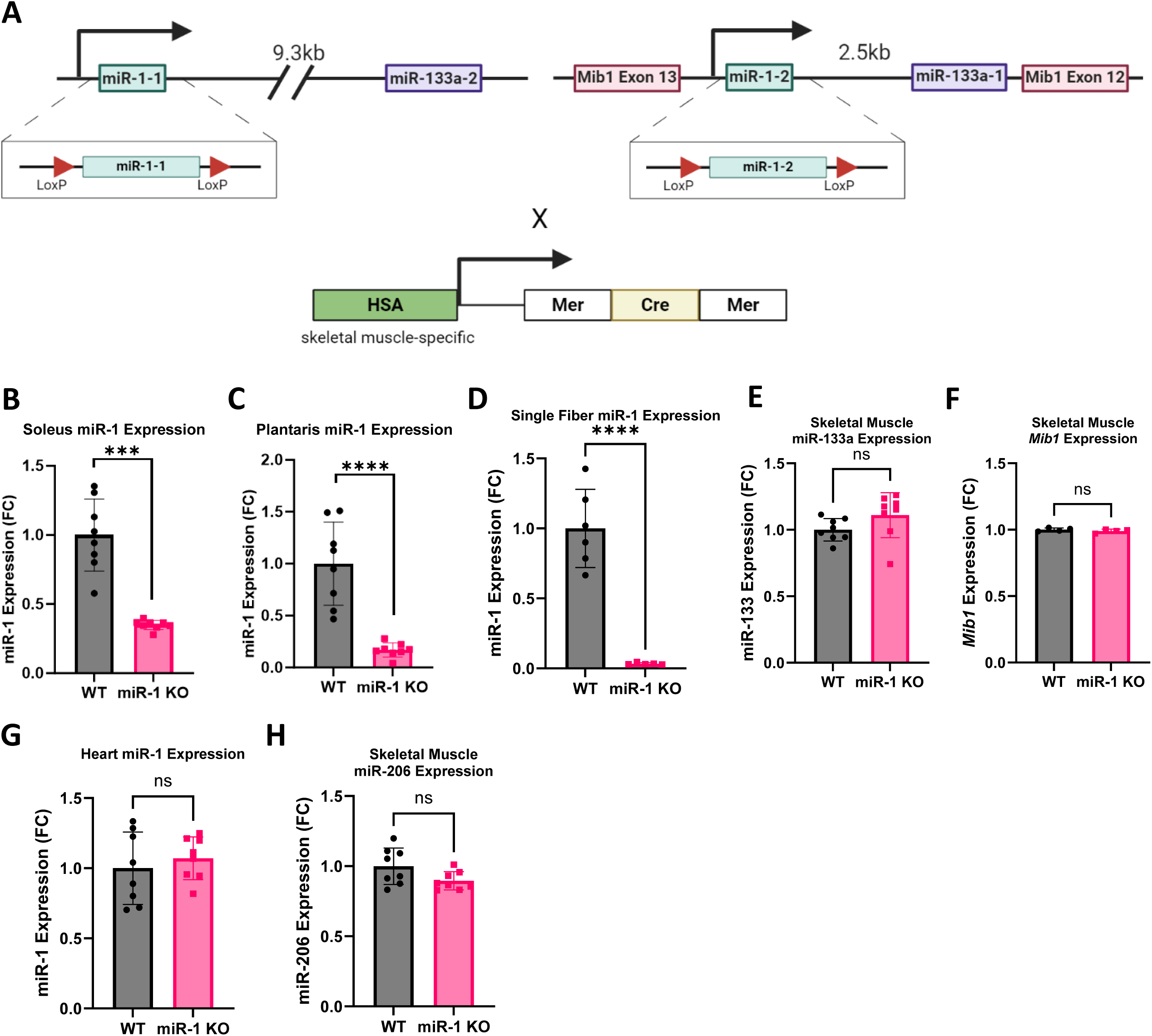
Generation of inducible, skeletal muscle-specific miR-1 KO mouse. **(A)** Targeting strategies to insert LoxP sites flanking genomic regions encompassing the *miR-1-1* and *miR-1-2* precursor to generate floxed alleles. Inducible, skeletal muscle-specific KO of miR-1 was achieved by breeding homozygous miR-1-1^f/f^; miR-1-2^f/f^ mice to a myofiber-specific CreER transgenic mouse in which the human alpha-skeletal muscle actin (HSA) promoter drives expression of CreER, allowing for recombination following tamoxifen administration. Created with Biorender.com. (**B-C)** Whole-muscle miR-1 expression in WT and miR-1 KO mice in **(B)** soleus and **(C)** plantaris muscles. **(D)** Single fiber miR-1 expression in WT and miR-1 KO mice. **(E)** Whole-muscle miR-133 expression and **(F)** *Mib1* mRNA expression**. (G)** miR-1 expression in heart tissue. **(H)** Whole-muscle miR-206 expression. FC: fold change, ns: not significant, \*\*\**p* < 0.001, \*\*\*\**p* < 0.0001. Independent *t*-tests used for comparisons. N=4-8 female mice per group (post-tamoxifen treatment and an 8-week washout).

**Supplemental Figure S2.**
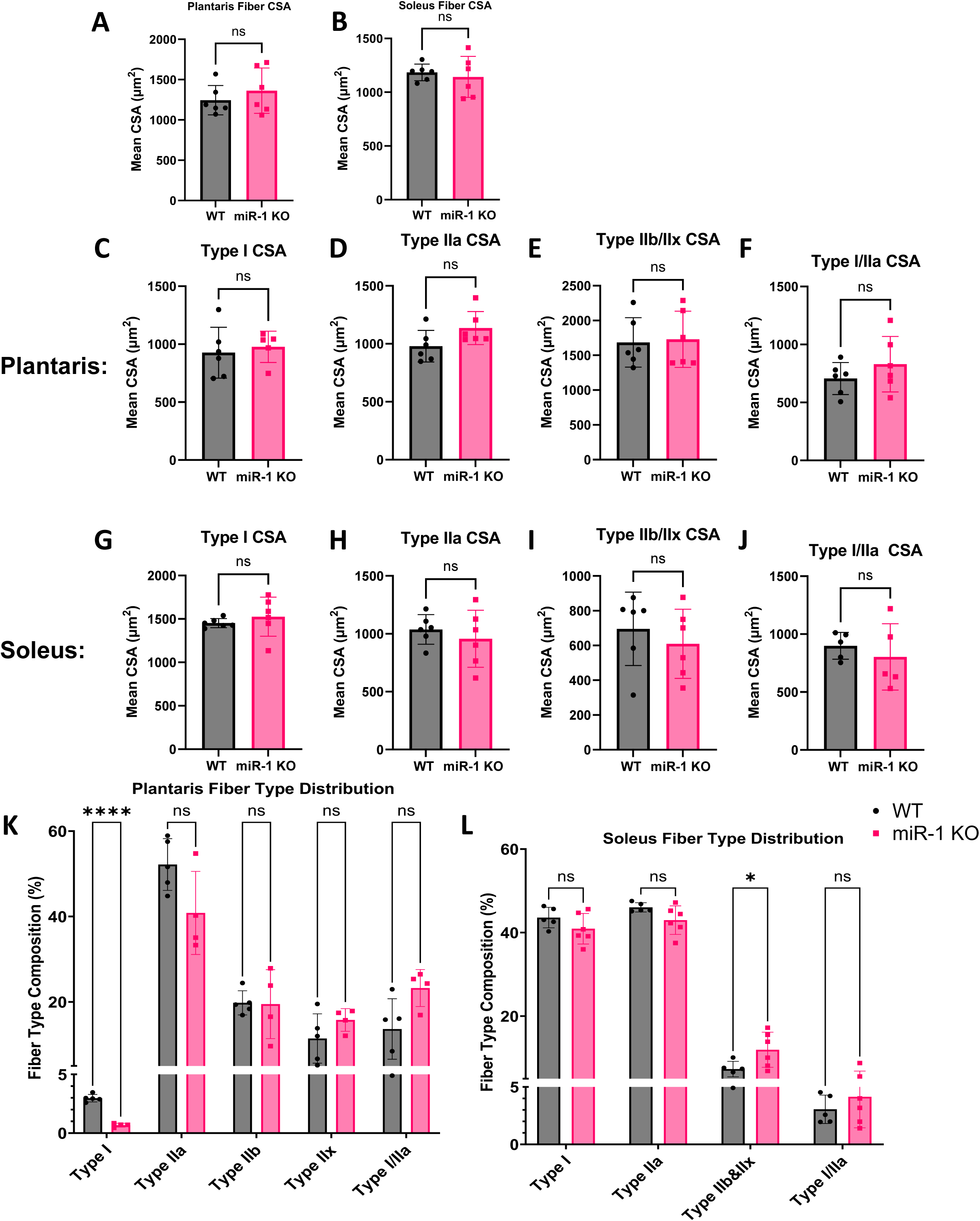
Muscle fiber characteristics of miR-1 KO mice. (A-B) Mean fiber CSA of **(A)** plantaris and **(B)** soleus muscles. **(C-F)** Fiber type-specific CSA of plantaris muscles, **(C)** Mean Type I fiber CSA, **(D)** Mean Type IIa fiber CSA, **(E)** Mean Type IIb and IIx fiber CSA, and **(F)** Mean hybrid Type I/IIa fiber CSA. **(G-J)** Fiber type-specific CSA of soleus muscles. **(G)** Mean Type I fiber CSA, **(H)** Mean Type IIa fiber CSA, **(I)** Mean Type IIb and IIx fiber CSA, and **(J)** Mean hybrid Type I/IIa fiber CSA. **(K-L)** Fiber type composition of **(K)** plantaris muscle and **(L)** soleus muscle. \**p* < 0.05, \*\*\*\**p* < 0.0001, ns: not significant. Independent *t*-tests used for comparisons, n=4-6 female mice per group (post-tamoxifen treatment and an 8-week washout).

**Supplemental Figure S3.**
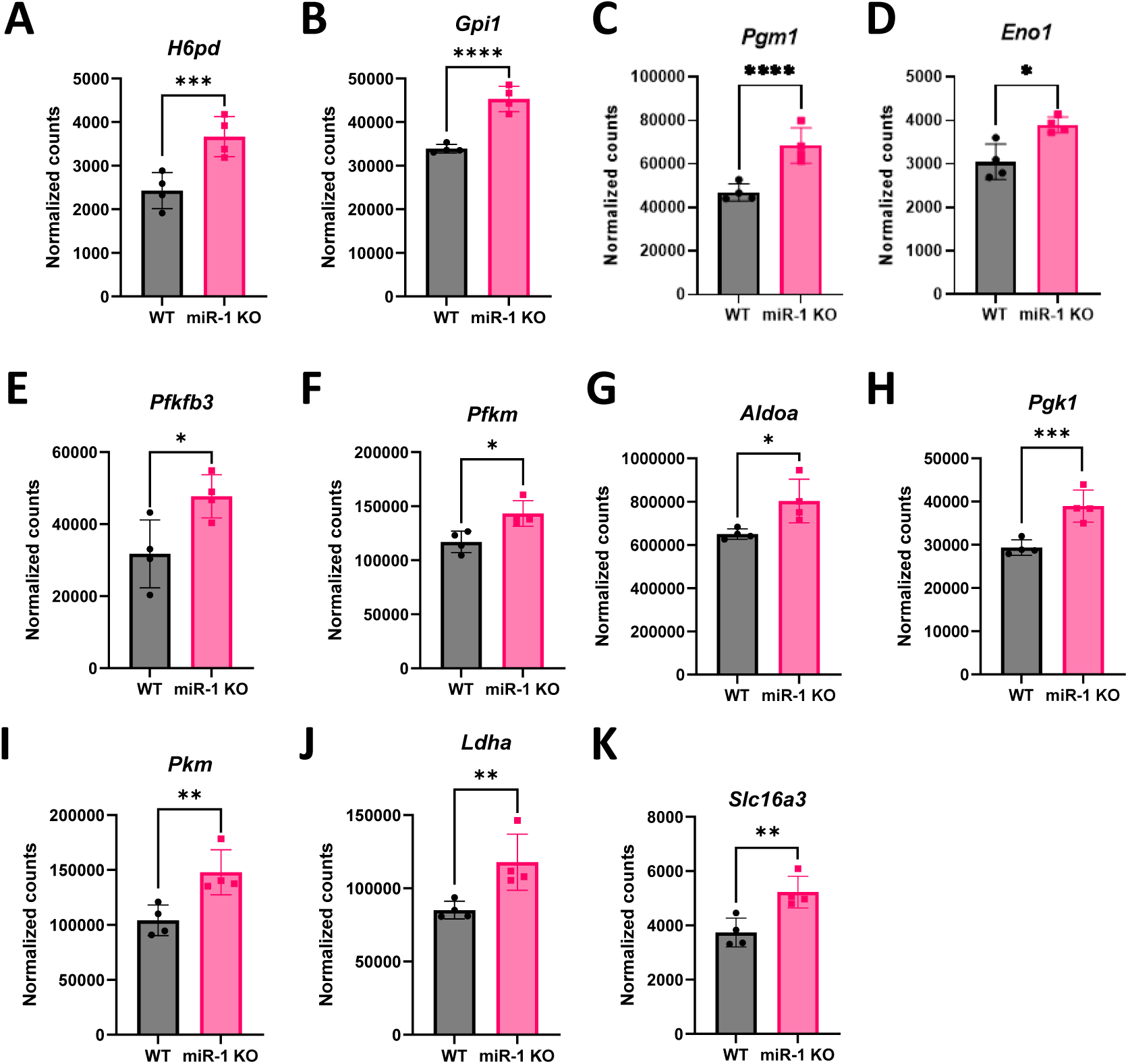
miR-1 regulation of glycolytic and pentose phosphate pathway gene expression. (A-K) Normalized RNA counts based on RNA-seq (n=4 female WT, n=4 female miR-1 KO, post-tamoxifen treatment and an 8-week washout) of **(A)** Hexose-6-phosphate dehydrogenase (*H6pd)*, **(B)** Glucose-6-phosphate isomerase 1 (*Gpi1*), **(C)** Phosphoglucomutase 1 (*Pgm1*), **(D)** Enolase 1 (*Eno1*), **(E)** 6-phosphofructo-2-kinase 3 (*Pfkfb3*), **(F)** Phosphofructokinase-muscle (*Pfkm*), **(G)** Aldolase A (*Aldoa*), **(H)** Phosphoglycerate Kinase 1 (*Pgk1*), **(I)** Pyruvate kinase muscle (*Pkm*), **(J)** Lactate dehydrogenase A (*Ldha*), and **(K)** Solute carrier family 16 member 3 (*Slc16a3*). FDR values indicated as \**p* < 0.05, \*\**p* < 0.01, \*\*\**p* < 0.001, \*\*\*\**p* < 0.0001.

**Supplemental Figure S4.**
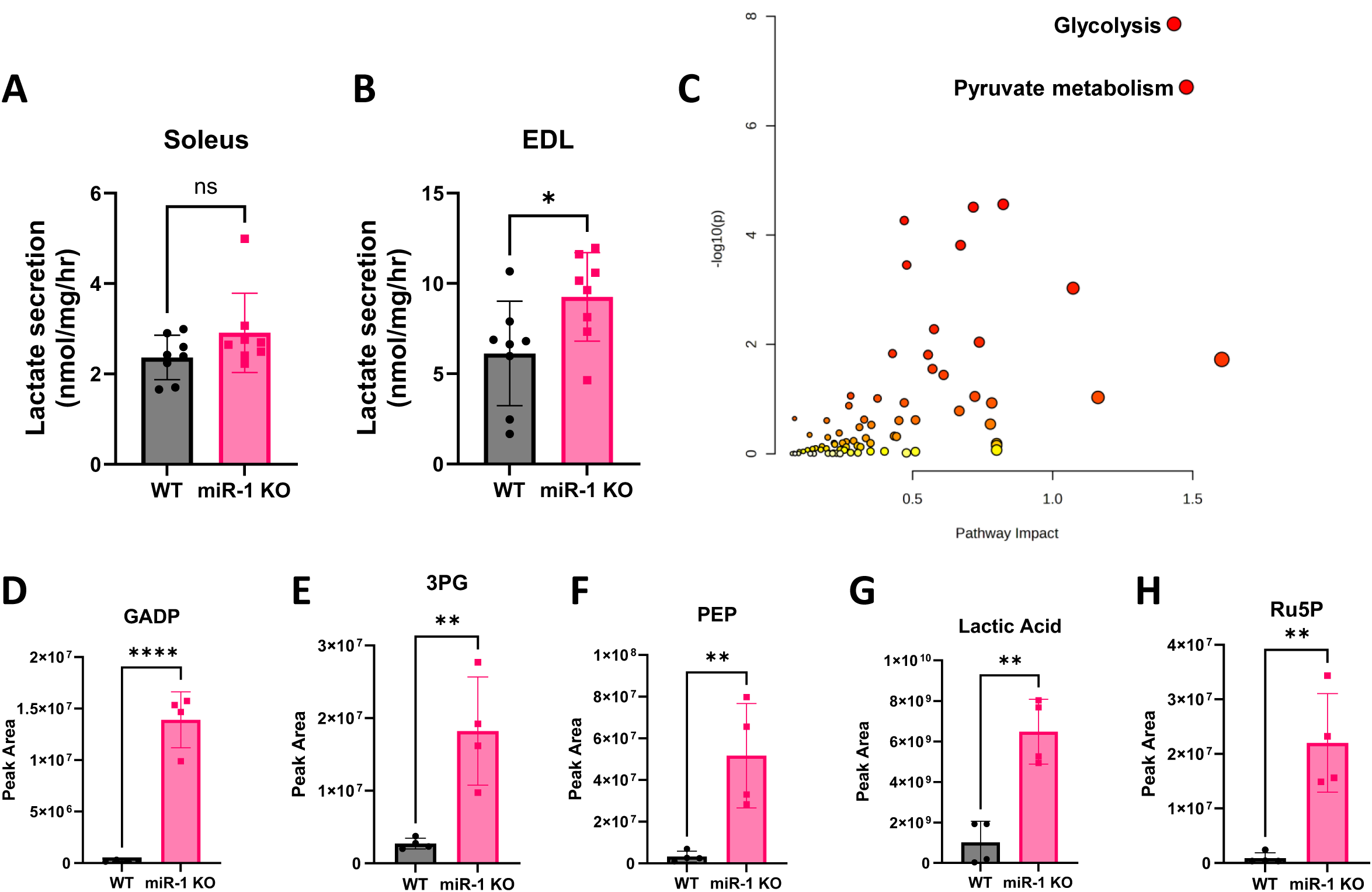
Differentially expressed glycolytic metabolites in miR-1 KO mice. (A-B) Buffer lactate levels measured as an indicator of lactate secretion of **(A)** soleus muscles and **(B)** EDL muscles, n= 8 female mice per group (post-tamoxifen treatment and an 8-week washout). **(C)** Summary of integrative analysis of miR-1 KO transcriptomics and metabolomics data at the pathway level, reflecting the impact on the pathway and the level of significance. **(D-H)** Peak Area of select glycolytic metabolites from metabolomics analyses of n=4 female WT and n=4 female miR-1 KO gastrocnemius muscles (post-tamoxifen treatment and an 8-week washout). **(D)** Glyceraldehyde 3-phosphate: GADP, **(E)** 3-phosphoglycerate: 3PG, **(F)** phosphoenolpyruvate: PEP, **(G)** Lactic acid, and **(H)** Ribulose 5-phosphate: Ru5P. Independent *t*-tests used for comparisons, ns: not significant, \**p* < 0.05, \*\**p* < 0.01, \*\*\*\**p* < 0.0001.

**Supplemental Figure S5.**
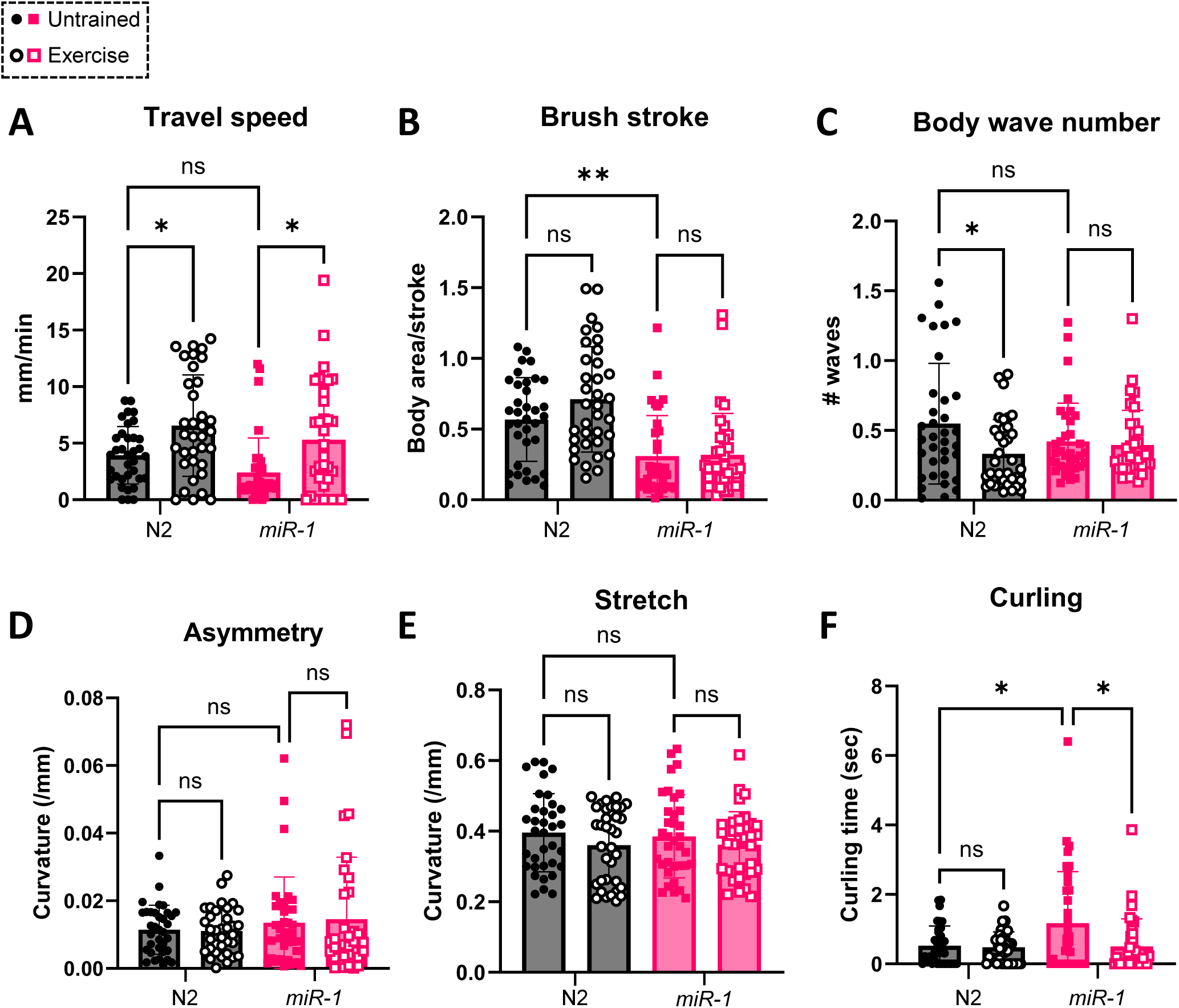
Physiological activity in *mir-1* mutant *C. elegans.* CeLeST analysis of additional physiological activity measures: travel speed, brush stroke, body wave number, asymmetry, stretch, and curling in Exercised (open circle/square) and Untrained (dark circle/square) N2 and *mir-1* mutant worms. Groups compared using independent *t*-tests, ns: not significant, \**p* < 0.05, \*\**p* < 0.01.

## Data Availability

Further information and requests for resources and reagents should be directed to and will be fulfilled by the Lead Contact (J.J.M.). AGO2 eCLIP-seq data are available on UCSC Genome Browser (https://genome.ucsc.edu/s/bdpeck/Ago2%20eCLIP). Bulk RNA-seq data have been deposited in the Gene Expression Omnibus (GEO) database (GSE274378).

## Acknowledgements

We thank Marsha Ensor and the University of Kentucky Energy Balance and Body Composition Core for the calorimetry expertise and the University of Kentucky Mass Spectrometry and Proteomics Core for the metabolomic expertise. We thank Djamel Lebeche, PhD, formerly of the Cardiovascular Institute, Icahn School of Medicine at Mount Sinai and currently at the University of Tennessee Health Science Center, for helping us re-derive the miR-1 floxed mice.

## Funding

This work was supported by National Institutes of Health grants from the National Institute on Aging (R01AG069909 to C.A.P. and J.J.M. and R00AG063994 to K.A.M.) and the National Heart, Lung and Blood Institute (HL144717 and HL150557 to R.L.B.).

## Disclosures

Y.W. is the owner of MyoAnalytics, LLC. The other authors have no conflict of interests to report.

## Notes

### Competing Interest Statement

The authors have declared no competing interest.

